# A glycosylphosphatidylinositol-anchored α-amylase encoded by *amyD* contributes to a decrease in the molecular mass of cell wall α-1,3-glucan in *Aspergillus nidulans*

**DOI:** 10.1101/2021.11.24.469953

**Authors:** Ken Miyazawa, Takaaki Yamashita, Ayumu Takeuchi, Yuka Kamachi, Akira Yoshimi, Yuto Tashiro, Ami Koizumi, Shigekazu Yano, Shin Kasahara, Motoaki Sano, Youhei Yamagata, Tasuku Nakajima, Keietsu Abe

## Abstract

α-1,3-Glucan is one of the main polysaccharides in the cell wall of *Aspergillus nidulans*. We previously revealed that it plays a role in hyphal aggregation in liquid culture, and that its molecular mass (MM) in an *agsA*-overexpressing (*agsA^OE^*) strain was larger than that in an *agsB*-overexpressing (*agsB^OE^*) strain. The mechanism that regulates the MM of α-1,3-glucan is poorly understood. Although the gene *amyD*, which encodes glycosyl-phosphatidylinositol (GPI)-anchored α-amylase (AmyD), is involved in the biosynthesis of α-1,3-glucan in *A. nidulans*, how it regulates this biosynthesis remains unclear. Here we constructed strains with disrupted *amyD* (Δ*amyD*) or overexpressed *amyD* (*amyD^OE^*) in the genetic background of the ABPU1 (wild-type), *agsA^OE^*, or *agsB^OE^* strain, and characterized the chemical structure of α-1,3-glucans in the cell wall of each strain, focusing on their MM. The MM of α-1,3-glucan from the *agsB^OE^ amyD^OE^* strain was smaller than that in the parental *agsB^OE^* strain. In addition, the MM of α-1,3-glucan from the *agsA^OE^* Δ*amyD* strain was greater than that in the *agsA^OE^* strain. These results suggest that AmyD is involved in decreasing the MM of α-1,3-glucan. We also found that the C-terminal GPI-anchoring region is important for these functions.

## Introduction

The fungal cell wall, composed mainly of polysaccharides, is essential for the survival of the fungus (Latgè et al., 2017). It has recently been understood that the cell wall is a highly dynamic structure; cell-wall components are synthesized by synthases and then reconstructed by glycosyltransferases to form a proper cell-wall architecture (Latgè and Beauvais, 2014;Latgè et al., 2017). The cell wall of filamentous fungi contains α-glucans, β-glucans, chitin, and galactomannan. Some fungi form an extracellular matrix, which includes secretory polysaccharides such as galactosaminogalactan (Sheppard and Howell, 2016;Yoshimi et al., 2016;Miyazawa et al., 2019). Cell-wall polysaccharides of some *Aspergillus* species can be fractionated into alkali-soluble and alkali-insoluble fractions (Fontaine et al., 2000;Yoshimi et al., 2013;Dichtl et al., 2015;Zhang et al., 2017b). The alkali-soluble fraction contains mainly α-1,3-glucan with interconnecting α-1,4-linkage and some galactomannan (Bernard and Latge, 2001;Latgè, 2010). The alkali-insoluble fraction is composed of chitin, β-1,6-branched β-1,3-glucan, and galactomannan (Fontaine et al., 2000;Bernard and Latge, 2001).

In the human pathogenic dimorphic yeast *Histoplasma capsulatum* and the rice blast fungus *Magnaporthe grisea*, α-1,3-glucan functions as a stealth factor that prevents host immune recognition and consequently contributes to the establishment of invasion or infection (Rappleye et al., 2004;Rappleye et al., 2007;Fujikawa et al., 2009;Fujikawa et al., 2012). In addition, the pathogenesis of an α-1,3-glucan-deficient strain is decreased in murine models infected with *Aspergillus fumigatus* (Henry et al., 2012;Beauvais et al., 2013). Recently, α-1,3-glucan was reported to stimulate the polarization of regulatory T-cells by inducing programmed death-ligand 1 expression on human dendritic cells (Stephen-Victor et al., 2017). Fontaine et al. (2010) revealed that α-1,3-glucan has adhesivity when the conidia of *A*. *fumigatus* germinate.

Grün et al. (2005) analyzed the detailed chemical structure of α-glucan in the cell wall of the fission yeast *Schizosaccharomyces pombe* and found that its molecular mass (MM) is 42 □ 600 ± 5 □200, which is equivalent to a degree of polymerization of 263 ± 32 (Grün et al., 2005). The α-glucans derived from *S. pombe* are composed of two chains of ≈120 residues of 1,3-linked α-glucose with 12 residues of 1,4-linked α-glucose at the reducing ends (Grün et al., 2005). In *Aspergillus wentii*, the water-insoluble (alkali-soluble) glucan has a MM of 850 000 and consists of 25 subunits (200 residues each) of α-1,3-glucan separated by short spacers composed of 1,4-linked α-glucan (Choma et al., 2013).

*Aspergillus* species have several α-1,3-glucan synthase genes: two in *Aspergillus nidulans* (*agsA* and *agsB*), three in *A*. *fumigatus* (*AGS1–3*) and *Aspergillus oryzae* (*agsA–C*), and five in *Aspergillus niger* (*agsA–E*). Disruptants of *A. fumigatus* that lack a single gene or all three genes have been constructed (Beauvais et al., 2005;Maubon et al., 2006;Henry et al., 2012); these strains lack α-1,3-glucan in the cell wall and are less pathogenic (Beauvais et al., 2013). In *A*. *oryzae, agsB* (orthologous to *A*. *nidulans agsB*) is the primary α-1,3-glucan synthase gene (Zhang et al., 2017b). An *A*. *oryzae* disruptant lacking all three genes loses its cell-wall α-1,3-glucan and forms small hyphal pellets under liquid culture conditions (Miyazawa et al., 2016). In *A*. *niger*, the expression of *agsA* (orthologous to *A*. *fumigatus AGS3*; no orthologue in *A*. *nidulans*) and *agsE* (orthologous to *A*. *nidulans agsB*) is upregulated in the presence of stress-inducing compounds in the cell wall (Damveld et al., 2005). In the kuro (black) koji mold *Aspergillus luchuensis*, disruption of *agsE* (orthologous to *A*. *nidulans agsB*) improves the protoplast formation (Tokashiki et al., 2019). Recently Uechi et al. revealed that *A*. *luchuensis agsB* (no orthologue in *A*. *nidulans*) plays a role in nigeran synthesis (Uechi et al., 2021). In *A*. *nidulans*, α-1,3-glucan in vegetative hyphae is synthesized mainly by AgsB (Yoshimi et al., 2013;He et al., 2014). The hyphae of a mutant deficient in α-1,3-glucan became fully dispersed, showing that α-1,3-glucan is a hyphal aggregation factor (Yoshimi et al., 2013;He et al., 2014). We recently constructed strains overexpressing *agsA* (*agsA^OE^*) and *agsB* (*agsB^OE^*) in the genetic background of, respectively, *agsB* and *agsA* disruptants. The peak MM of alkali-soluble glucan from *agsA^OE^* was 1 480 000 ± 80 000, which was four times that from the *agsB^OE^* (MM, 372 □ 000 ± 47≥ 000) (Miyazawa et al., 2018). The alkali-soluble glucan derived from these strains contains several 1,4-linked spacer structures interlinking the α-1,3-glucan subunits, which each contain 200 glucose residues (Miyazawa et al., 2018).

Outside of *A*. *fumigatus*, *A. nidulans agsB* and its orthologues are clustered with two α-amylase-encoding genes (*amyD* and *amyG* in *A*. *nidulans*) (He et al., 2014;Yoshimi et al., 2017;Miyazawa et al., 2020). The *amyG* gene encodes an intracellular α-amylase and is crucial for α-1,3-glucan synthesis (He et al., 2014). The *amyD* gene in *A*. *nidulans* encodes glycosylphosphatidylinositol (GPI)-anchored α-amylase. He et al. (2014) reported that α-1,3-glucan contents increased by 50% in an *amyD*-disrupted (Δ*amyD*) strain and halved in an *amyD*-overexpressing (*actA*(p)-*amyD*) strain, suggesting that *amyD* has a repressive effect on α-1,3-glucan synthesis. In addition, He et al. (2017) analyzed the chronological changes of α-1,3-glucan contents under liquid culture conditions. Whereas the amount of α-1,3-glucan in strains that overexpressed the α-1,3-glucanase-encoding gene (*mutA* or *agnB*) was decreased after 20 h from inoculation, the amount of α-1,3-glucan in the cell wall of the *amyD^OE^* strain was half that of the wild-type strain from the initial stage of cultivation (He et al., 2017). He et al. (2017) suggested that AmyD decreased the amount of α-1,3-glucan in the cell wall by a mechanism independent of the effect of α-1,3-glucanase. The enzymatic characteristics of *A*. *niger* AgtA, which is encoded by an orthologue of *A*. *nidulans amyD*, have been reported (van der Kaaij et al., 2007). Although AgtA in *A. niger* barely hydrolyzed α-1,3-glucan, it had relatively high transglycosylation activity on donor substrates with maltooligosaccharides (van der Kaaij et al., 2007). Overall, AmyD seems to indirectly decrease the amount of α-1,3-glucan in the cell wall, but the detailed mechanism is still unknown.

Here, in a study of the function of *amyD* in α-1,3-glucan biosynthesis in *A. nidulans*, we constructed strains with overexpression or disruption of *amyD* in the genetic backgrounds of the wild-type, *agsA^OE^*, and *agsB^OE^*. We performed several chemical analyses of α-1,3-glucan derived from the strains, looking in particular at its MM, and examined the role of *amyD* in controlling the MM of α-1,3-glucan in the cell wall.

## Materials and Methods

### Strains and growth media

Strains are listed in Table 1. Czapek-Dox (CD) medium was used as the standard culture, as described previously (Fujioka et al., 2007;Miyazawa et al., 2018).

**Table 1.** Strains used in this study.

| Strains | Genotype | References |
| --- | --- | --- |
| A4 |  | FGSC<sup>a</sup> |
| ABPU1 (*argB*<sup>+</sup>)<br>(wild-type) | *biA1, pyrG89, wA3, argB2, pyroA4, veA1, ligD::ptrA, AoargB*<sup>+</sup> | (Hagiwara et al., 2007; Miyazawa et al., 2018) |
| $\Delta amyD$ | *biA1, pyrG89, wA3, argB2, pyroA4, veA1, ligD::ptrA, AoargB*<sup>+</sup>, *amyD::pyrG* | This study |
| *amyD*<sup>OE</sup> | *biA1, pyrG89, wA3, argB2, pyroA4, veA1, ligD::ptrA, AoargB*<sup>+</sup>, *PtefI-amyD::pyrG* | This study |
| $\Delta agsA$ | *biA1, pyrG89, wA3, argB2, pyroA4, veA1, ligD::ptrA, AoargB*<sup>+</sup>, *agsA::loxP* | This study |
| *agsB*<sup>OE</sup> | *biA1, pyrG89, wA3, argB2, pyroA4, veA1, ligD::ptrA, AoargB*<sup>+</sup>, *agsA::loxP*, *PtefI-agsB::pyroA* | This study |
| *agsB*<sup>OE</sup> $\Delta amyD$ | *biA1, pyrG89, wA3, argB2, pyroA4, veA1, ligD::ptrA, AoargB*<sup>+</sup>, *agsA::loxP*, *PtefI-agsB::pyroA, amyD::pyrG* | This study |
| *agsB*<sup>OE</sup> *amyD*<sup>OE</sup> | *biA1, pyrG89, wA3, argB2, pyroA4, veA1, ligD::ptrA, AoargB*<sup>+</sup>, *agsA::loxP*, *PtefI-agsB::pyroA, PtefI-amyD::pyrG* | This study |
| $\Delta agsB$ | *biA1, pyrG89, wA3, argB2, pyroA4, veA1, ligD::ptrA, agsB::argB* | (Yoshimi et al., 2013) |
| *agsA*<sup>OE</sup> | *biA1, pyrG89, wA3, argB2, pyroA4, veA1, ligD::ptrA, agsB::argB, PtefI-agsA::pyroA* | This study |
| *agsA*<sup>OE</sup> $\Delta amyD$ | *biA1, pyrG89, wA3, argB2, pyroA4, veA1, ligD::ptrA, agsB::argB, PtefI-agsA::pyroA, amyD::pyrG* | This study |
| *agsA*<sup>OE</sup> *amyD*<sup>OE</sup> | *biA1, pyrG89, wA3, argB2, pyroA4, veA1, ligD::ptrA, agsB::argB, PtefI-agsA::pyroA, PtefI-amyD::pyrG* | This study |
| $\Delta amyD$-*amyD*<sup>OE</sup> | *biA1, pyrG89, wA3, argB2, pyroA4, veA1, ligD::ptrA, AoargB*<sup>+</sup>, *amyD::pyrG, PtefI-amyD::hph, pyrG*<sup>-</sup> | This study |
| $\Delta amyD$-*amyD*<sup>OE</sup>($\Delta GPI$) | *biA1, pyrG89, wA3, argB2, pyroA4, veA1, ligD::ptrA, AoargB*<sup>+</sup>, *amyD::pyrG, PtefI-amyD($\Delta GPI$)::hph, pyrG*<sup>-</sup> | This study |
| *agsB*<sup>OE</sup> $\Delta amyD$-*amyD*<sup>OE</sup> | *biA1, pyrG89, wA3, argB2, pyroA4, veA1, ligD::ptrA, agsB::argB, PtefI-agsA::pyroA, amyD::pyrG, PtefI-amyD::hph, pyrG*<sup>-</sup> | This study |
| *agsB*<sup>OE</sup> $\Delta amyD$-*amyD*<sup>OE</sup>($\Delta GPI$) | *biA1, pyrG89, wA3, argB2, pyroA4, veA1, ligD::ptrA, agsB::argB, PtefI-agsA::pyroA, amyD::pyrG, PtefI-amyD($\Delta GPI$)::hph, pyrG*<sup>-</sup> | This study |

<sup>a</sup> Fungal Genetic Stock Center, USA

### Construction of the *agsA*- and *agsB*-overexpressing strains

We newly constructed *agsA^OE^* and *agsB^OE^* strains for this study. To generate *agsA^OE^*, pAPyT-agsA plasmids (Miyazawa et al., 2018) were digested with *NotI* and transformed into a disrupted *agsB* (Δ*agsB*) strain (Fig. S1A). Correct integration of *agsA* overexpression cassettes was confirmed by PCR (Fig. S1B). To generate *agsB^OE^*, the disrupted *agsA* (Δ*agsA*) strain was first generated using the Cre/*loxP* marker recycling system (Zhang et al., 2017a). The pAPG-cre/DagsA plasmid (Miyazawa et al., 2018) was digested with *Eco*RI and transformed into the ABPU1 (*argB*^+^) strain. Candidate strains were selected on CD medium without uridine and uracil, and then cultured on CD medium with uridine and uracil and 1% xylose to induce *Cre* expression (Fig. S1C). Strains that required uridine and uracil were isolated, and then replacement of the *agsA* gene was confirmed by PCR (Fig. S1D). The pAPyT-agsB plasmid was digested with *NotI* and transformed into the Δ*agsA* strain (Fig. S1E). Correct integration of *agsB* overexpression cassettes was confirmed by PCR (Fig. S1B).

### Construction of the *amyD^OE^* strain

The *amyD^OE^* strain was constructed by replacing the native promoter with the constitutive *tef1* promoter. The sequences of the primers are listed in Table S1. To generate *amyD^OE^*, the plasmid pAPT-amyD was constructed (Fig. S2A). The 5’-non-coding region (amplicon 1) and the coding region (amplicon 2) of *amyD* were amplified from *A*. *nidulans* ABPU1 genomic DNA. The *pyrG* marker (amplicon 3) was amplified from the pAPG-cre/DagsA plasmid. The *tef1* promoter (amplicon 4) was amplified from the pAPyT-agsB plasmid. The four amplicons and a *Sac*I-digested pUC19 vector were fused using an In-Fusion HD Cloning Kit (Clontech Laboratories, Inc., Mountain View, CA, USA). The resulting plasmid was digested with *SacI*, and transformed into the ABPU1 (*argB*^+^), *agsA^OE^*, and *agsB^OE^* strains (Fig. S2B). Correct integration of the cassette was confirmed by PCR (Fig. S2C).

### Disruption of the *amyD* gene

In the first round of PCR, gene fragments containing the 5’-non-coding region (amplicon 1) and the coding region (amplicon 2) of *amyD* were amplified from ABPU1 genomic DNA, and the *pyrG* gene (amplicon 3) was amplified from *A*. *oryzae* genomic DNA (Fig. S2D). The three resulting fragments were gel-purified and fused into a disruption cassette in the second round of PCR. The resulting PCR product was gel-purified and transformed into the ABPU1 (*argB*^+^), *agsA^OE^*, and *agsB^OE^* strains (Fig. S2E). Replacement of the *amyD* gene was confirmed by PCR (Fig. S2F).

### Expression of complementary *amyD* genes

The sequences of the primers are listed in Table S1. A GPI-anchor modification site, the *ω*-site, was predicted with the GPI Prediction Server v. 3.0 (https://mendel.imp.ac.at/gpi/gpi_server.html), and the best score for the ω-site was Asn535 of AmyD. To remove the GPI anchor of AmyD, 54 nucleotides corresponding to the 18 amino acid residues from Asn535 in AmyD were deleted from the authentic *amyD* gene (Fig. S3A). To create complementary genes that have full-length open reading frames of either *amyD* or the gene without the GPI anchor–coding region, the plasmids pAHT-amyD, pAHdPT-amyD, pAHT-amyD(ΔGPI), and pAHdPT-amyD(ΔGPI) were first constructed (Fig. S3A). To construct pAHT-amyD, primers IF-Ptef1-hph-Fw and IF-amyD-up-hph-Rv were amplified by PCR using pAPT-amyD as a template (amplicon 1). The hygromycin-resistance gene *hph* (amplicon 2) was amplified with primers 397-5 and 397-3 from pSK397 (Krappmann et al., 2006). The two amplicons were fused using a NEBuilder HiFi DNA Assembly kit (New England Biolabs, Ipswich, MA, USA) according to the manufacturer’s instructions. Then, to delete the GPI anchor–encoding region of *amyD*, PCR amplification was performed with primers ANamyD-dGPI-Fw and ANamyD-dGPI-Rv from the resulting pAHT-amyD plasmid with PrimeSTAR Max DNA Polymerase (Takara Bio Inc., Kusatsu, Japan). The amplified fragment was transformed into DH5α competent cells, and the pAHT-amyD(ΔGPI) plasmid was obtained (Fig. S3A). To construct pAHdPT-amyD, the first half (amplicon 1) and the second half (amplicon 2) of *pyrG* were amplified from *A*. *oryzae* genomic DNA. The fragment containing *hph*, *tef1* promoter, and *amyD* (amplicon 3) was amplified from pAHT-amyD. The three amplicons were fused using a NEBuilder kit. For pAHdPT-amyD(ΔGPI) construction, the fragment containing *hph*, *tef1* promoter, and *amyD* lacking its GPI anchor–coding region (amplicon 3’) was amplified from pAHT-amyD(ΔGPI). The three amplicons and the *Sac*I-digested pUC19 vector were fused using an In-Fusion HD Cloning Kit (Fig. S3B). The pAHdPT-amyD and pAHdPT-amyD(ΔGPI) plasmids were digested with *SacI* and transformed into the Δ*amyD* and *agsB^OE^* Δ*amyD* strains (Fig. S3C). Correct integration of the cassettes was confirmed by PCR (Fig. S3D).

### RNA extraction and quantitative real-time PCR

Mycelial cells cultured in CD liquid medium for 24 h were collected, and total RNA was extracted from the cells by using Sepasol-RNA I Super G (Nakalai Tesque, Kyoto, Japan) in accordance with the manufacturer’s instruction. The total RNA (2.5 μg) was reverse-transcribed by using a SuperScript IN VILO Master Mix with ezDNase Enzyme (Invitrogen, Carlsbad, CA, United States). Quantitative real-time PCR was performed with a Mx3000P (Agilent Technologies, Santa Clara, CA, United States) with SYBR Green detection. For reaction mixture preparation, Thunderbird Next SYBR qPCR Mix (Toyobo Co., Ltd., Osaka, Japan) was used. Primers used for quantitative PCR are listed in Table S1. An equivalent amount of cDNA, obtained from reverse transcription reactions using an equivalent amount of total RNA, was applied to each reaction mixture. The gene encoding histone H2B was used as a normalization reference (an internal control) for determining the target gene expression ratios.

### Delipidization and fractionation of mycelial cells

Cell walls were fractionated as previously described with some modification (Miyazawa et al., 2018). Mycelia cultured for 24 h in CD medium were collected by filtering through Miracloth (Merck Millipore, Darmstadt, Germany), washed with water, and freeze-dried. The mycelia were then pulverized in a MM400 bench-top mixer mill (Retch, Haan, Germany). The powder (1 g) was suspended in 25 mL of chloroform–methanol (3:1 vol/vol) and stirred at room temperature for 12 h to remove the total polar lipid content of the mycelial cells. The mixture was centrifuged (10 □ 000 × *g*, 10 min). The residue was suspended in chloroform– methanol, and the delipidizing procedure was repeated. Then the de-polar lipid residue was suspended in 40 mL of 0.1 M Na phosphate buffer (pH 7.0), and cell-wall components were fractionated by hot-water and alkali treatments, as described previously (Miyazawa et al., 2018). Hot-water–soluble, alkali-soluble, and alkali-insoluble fractions were obtained from this fractionation, and the alkali-soluble fraction was further separated into a fraction soluble in water at neutral pH (AS1) and an insoluble fraction (AS2). The monosaccharide composition of AS2 fractions was quantified according to Miyazawa et al. (2018).

To obtain mycelia cultured for 16 h, conidia (final conc. 5.0 × 10^5^/mL) were inoculated into 200 mL CD medium and rotated at 160 rpm at 37°C. The mycelia were collected and fractionated as described above.

### ^13^C NMR analysis

The AS2 fraction of each strain (50 mg) was suspended in 1 mL of 1 M NaOH/D_2_O and dissolved by vortexing. One drop of DMSO-d_6_ (deuterated dimethyl sulfoxide) was then added to each fraction and the solutions were centrifuged (3,000 × *g*, 5 min) to remove insoluble debris. ^13^C NMR spectra of the supernatants were obtained using a JNM-ECX400P spectrometer (JEOL, Tokyo, Japan) at 400 MHz at 35°C. Chemical shifts were recorded relative to the resonance of DMSO-d6.

### Determination of the average molecular mass of alkali-soluble glucan

The MM of alkali-soluble glucan was determined by gel permeation chromatography (GPC) according to the methods of Puanglek et al. (2016), with some modification. A GPC-101 system (Showa Denko Co. Ltd., Tokyo, Japan) with an ERC-3125S degasser (Showa Denko) and an RI-71S refractive index detector (Showa Denko) was used for the measurement. It was fitted with a GPC KD-G 4A guard column (Showa Denko) and a GPC KD-805 column (8.0 × 300 mm; Showa Denko). The eluent was 20 mM LiCl in *N*,*N*-dimethylacetamide (DMAc), and the flow rate was 0.6 mL/min at 40°C. The detector was normalized with polystyrene standards (SM-105; Showa Denko). With SmartChrom software (Jasco, Tokyo, Japan), the GPC profile was divided into virtual time slices (n_*i*_) with the height of each virtual slice from the base line (*H_i_*) corresponding to a certain MM (*M_i_*) obtained by calibrating the column. From these values, the number-average MM (*M*_n_) and weight-average MM (*M*_w_) were calculated as:

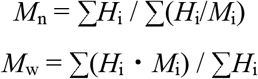

Polydispersity was calculated as *M*_w_/*M*_n_.

### Smith degradation

Smith degradation of the alkali-soluble glucan was performed as described (Miyazawa et al., 2018).

### Fluorescent labeling of cell-wall polysaccharides

Mycelial cells cultured for 16 h in CD liquid medium were dropped on a glass slide and dried at 55°C for 15 min. The cells were fixed, labeled with fluorophores, and imaged by confocal scanning microscopy as described (Miyazawa et al., 2018). Enzymatic digestion of β-1,3-glucan in the hyphal cells was performed as described (Miyazawa et al., 2018).

## Results

### Characterization of strains with disrupted or overexpressed *amyD*

We constructed *amyD^OE^* and Δ*amyD* strains by introducing the *amyD* cassettes for overexpression and disruption into the wild-type, *agsA^OE^*, and *agsB^OE^* strains (Fig. S2). The expression level of *amyD* in each strain was quantified in hyphal cells. Whereas each disrupted strain (Δ*amyD, agsA^OE^* Δ*amyD*, and *agsB^OE^* Δ*amyD*) showed scarce *amyD* expression, each overexpressing strain (*amyD^OE^*, *agsA^OE^ amyD^OE^*, and *agsB^OE^ amyD^OE^*) showed significantly higher *amyD* expression than the parental strains (Fig. 1).

**Figure 1.**
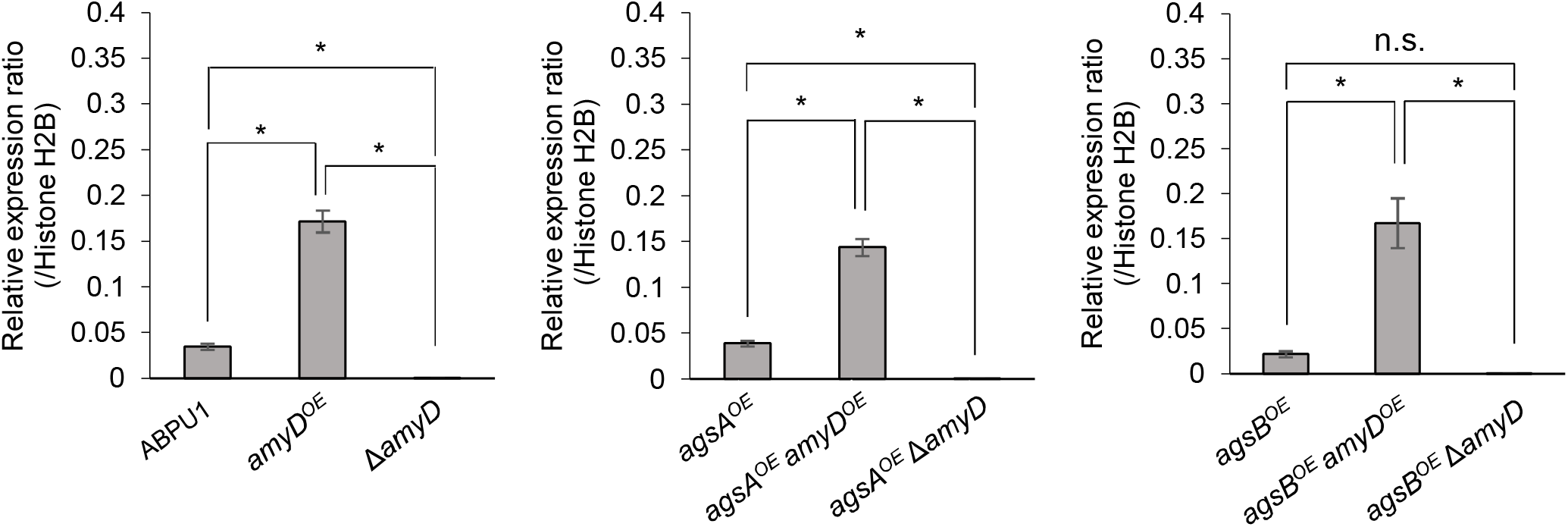
Transcript levels of the *amyD* gene determined by quantitative PCR. Genespecific primers are indicated in Table S1. Error bars represent the standard deviation of the mean calculated from three replicates. *Significant differences by Tukey’s test (*P* < 0.05); n.s., not significant.

There was no significant difference in radial growth among the strains grown on agar plates for 5 days (Fig. S4). In liquid culture, the wild-type and Δ*amyD* strains formed tightly aggregated hyphal pellets; however, the hyphae of the *amyD^OE^* strain were almost fully dispersed (Fig. 2). He et al. reported that the phenotype of their *amyD^OE^* strain resembles that of the Δ*agsB* strain in *A*. *nidulans* (He et al., 2014), which is consistent with our results (Fig. 2). In agreement with our previous results (Miyazawa et al., 2018), the *agsA^OE^* and *agsB^OE^* strains formed, respectively, loosely and tightly aggregated pellets (Fig. 2). Disruption of *amyD* did not affect the phenotypes of the *agsA^OE^* and *agsB^OE^* strains (Fig. 2). Also, overexpression of *amyD* scarcely affected the phenotypes of the *agsA^OE^* and *agsB^OE^* strains (Fig. 2).

**Figure 2.**
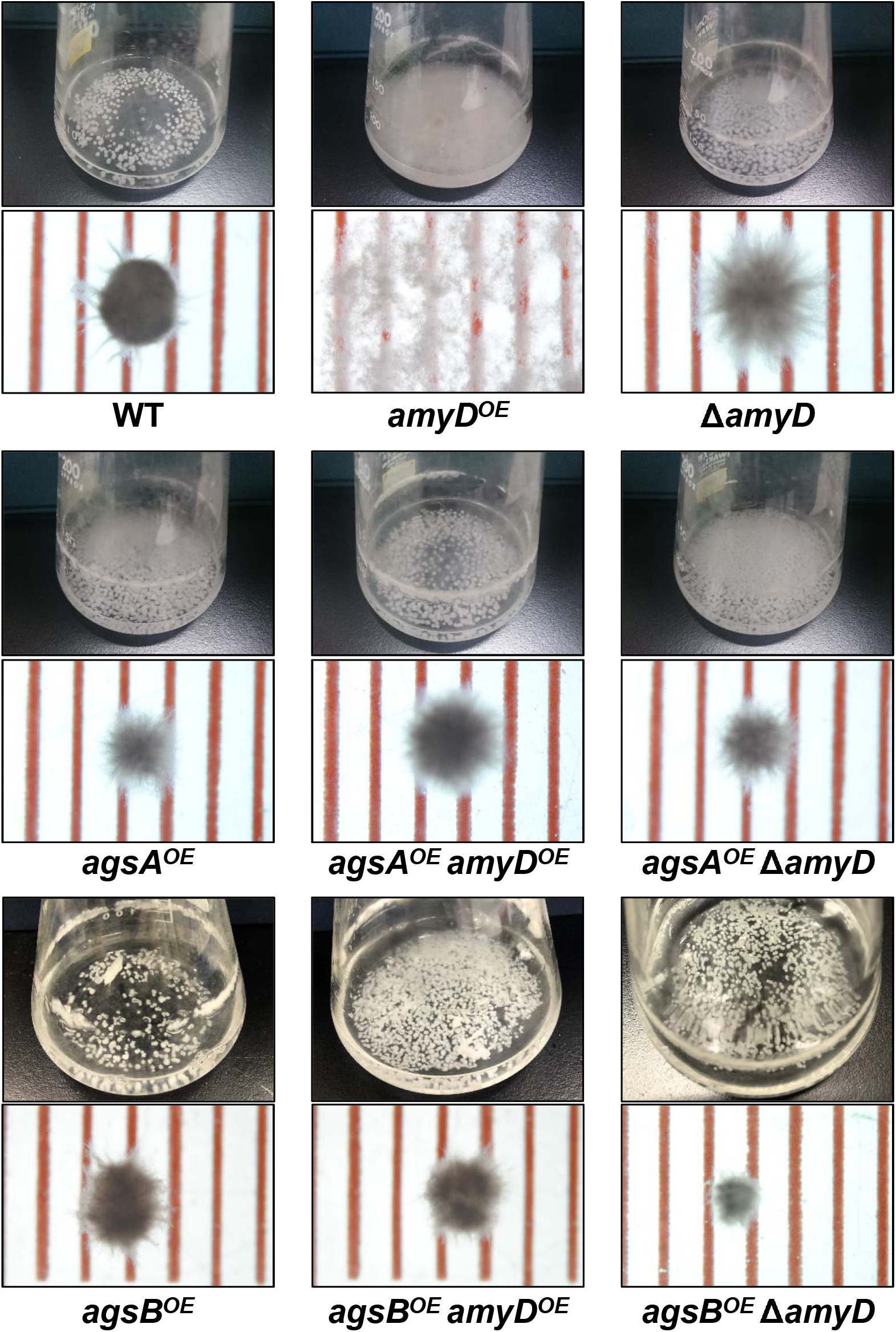
Growth characteristics of *amyD^OE^* and Δ*amyD* strains in liquid culture. Upper images, cultures in Erlenmeyer flasks; lower images, representative hyphal pellets of each strain under a stereomicroscope. Scale intervals are 1 mm.

### Overexpression of *amyD* resulted in a decrease in cell-wall alkali-soluble glucan

Cell-wall components of each strain were fractionated by a hot water–alkali treatment method, each fraction was weighed, and the monosaccharide composition of the AS2 fraction was quantified. The amount of glucose in the AS2 fraction was significantly lower in the *amyD^OE^* strain than in the wild-type strain (Fig. 3A; *P* < 0.05). That in the Δ*amyD* strain was similar to that in the wild-type strain (Fig. 3A). Those in the *agsA^OE^ amyD^OE^* and *agsA^OE^* Δ*amyD* strains, which were constructed from the parental strain *agsA^OE^*, were almost the same (Fig. 3B). It was significantly lower in the *agsB^OE^ amyD^OE^* strain than in the *agsB^OE^* and *agsB^OE^* Δ*amyD* strains (Fig. 3C; *P* < 0.05). These results indicate that AmyD acts to decrease the amount of alkali-soluble glucan in the wild-type and *agsB* strains, but not in the *agsA^OE^* strain, even when *amyD* is overexpressed.

**Figure 3.**
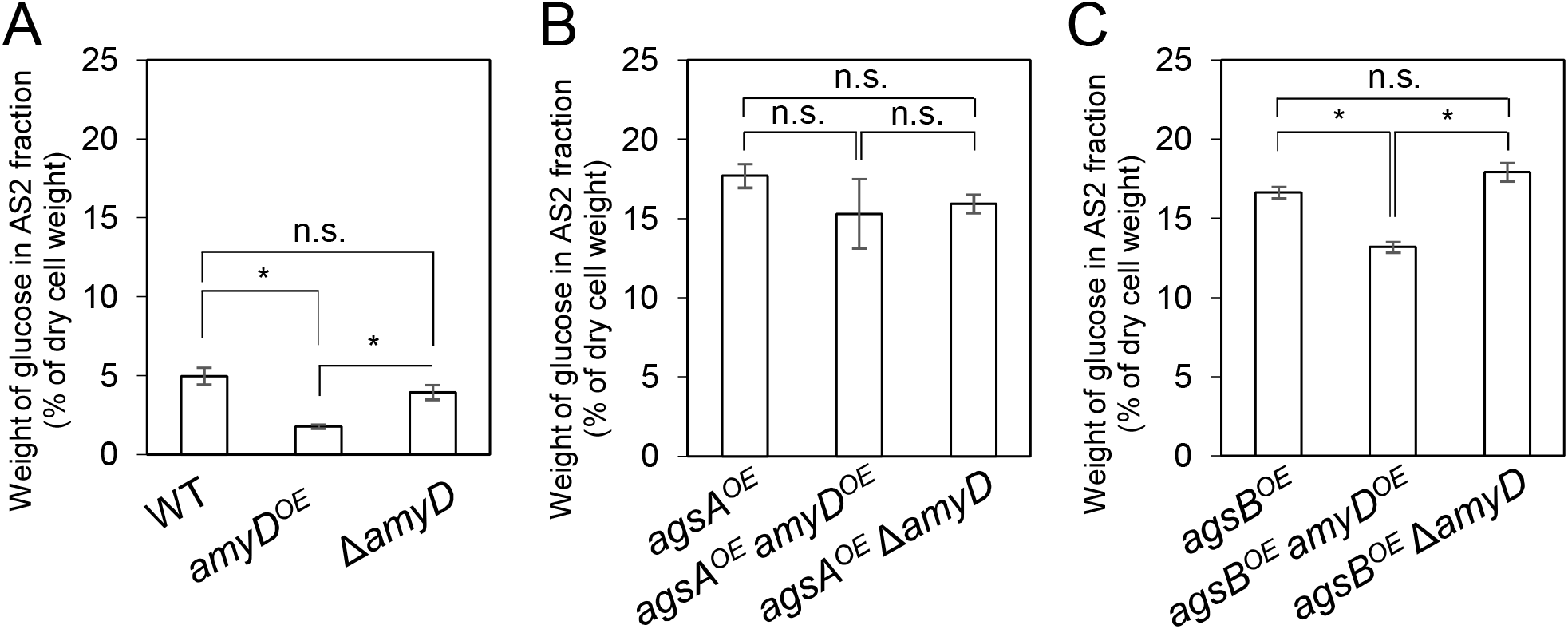
Amount of glucose in AS2 fractions. Conidia (5.0 × 10^5^/mL) of each strain were inoculated into CD medium and rotated at 160 rpm at 37°C for 24 h. Values show glucose content of the AS2 fraction as a percentage of the total cell-wall weight. Error bars represent standard error of the mean calculated from three replicates. *Significant difference by Tukey’s test (*P* < 0.05); n.s., not significant.

### Overexpression of the *amyD* gene decreases the molecular mass of alkali-soluble glucan

By ^13^C NMR analysis, the primary component in the AS2 fraction of the wild-type, *amyD^OE^*, and Δ*amyD* strains was found to be α-1,3-glucan, suggesting that *amyD* did not affect the primary components of alkali-soluble glucan (Fig. S5). To reveal whether the MM of alkali-soluble glucan was affected by disruption or overexpression of *amyD*, we determined the MM of alkali-soluble glucan in each strain by GPC analysis. Polystyrene (MM, 13□900– 3 ≥ 850 000) was used as a standard molecule to calibrate the column for size exclusion analysis. Although the physical properties of a polymer depend on *M*_w_, the number of moles is important for a biological reaction. Here we focus on the MM of alkali-soluble glucan with *M*_n_. The *M_n_* of the alkali-soluble glucan was 1 260 000 ± 270□000 in the *agsA^OE^* strain and 312 □ 000 ± 3□000 in *agsB^OE^* strain (Fig. 4A, B; Table 2), consistent with our previous results (Miyazawa et al., 2018). Although the *M*_n_ of alkali-soluble glucan in the *agsA^OE^ amyD^OE^* strain (1 □ 110 ≥ 000 ± 60 000) was similar to that in the parental (*agsA^OE^*) strain, that of *agsA^OE^ ΔamyD* was significantly greater (2 □ 250□000 ± 130 □ 000) than that of *agsA^OE^* (Fig. 4A; Table 2, *P* < 0.05). In addition, the *M_n_* of *agsB^OE^ amyD^OE^* (140 □ 000 ± 4 □ 000) was significantly less than that of the parental (*agsB^OE^*) strain (Fig. 4B; Table 2, *P* < 0.05). The *M*_n_ of alkali-soluble glucan in *agsB^OE^* Δ*amyD* (358 □ 000 ± 11 □ 000) was similar to that in *agsB^OE^* (Fig. 4B; Table 2). Lastly, the *M_n_* of alkali-soluble glucan in the wild type (2 □ 280 □ 000 ± 130□000) and Δ*amyD* (2□390□000 ± 390□000) was larger than that in *agsB^OE^* (312□000 ± 3□000; Fig. 4C; Table 2). The *amyD^OE^* strain had a primary peak at around 17 min (*M*_n_^1^, 32□900 ± 300) and a secondary peak at 11 min (*M*_n_^2^, 2□210□000 ± 700□000). These results suggest that AmyD degraded the alkali-soluble glucan eluted around 11 min to produce alkali-soluble glucan with a smaller MM (Fig. 4C; Table 2).

**Figure 4.**
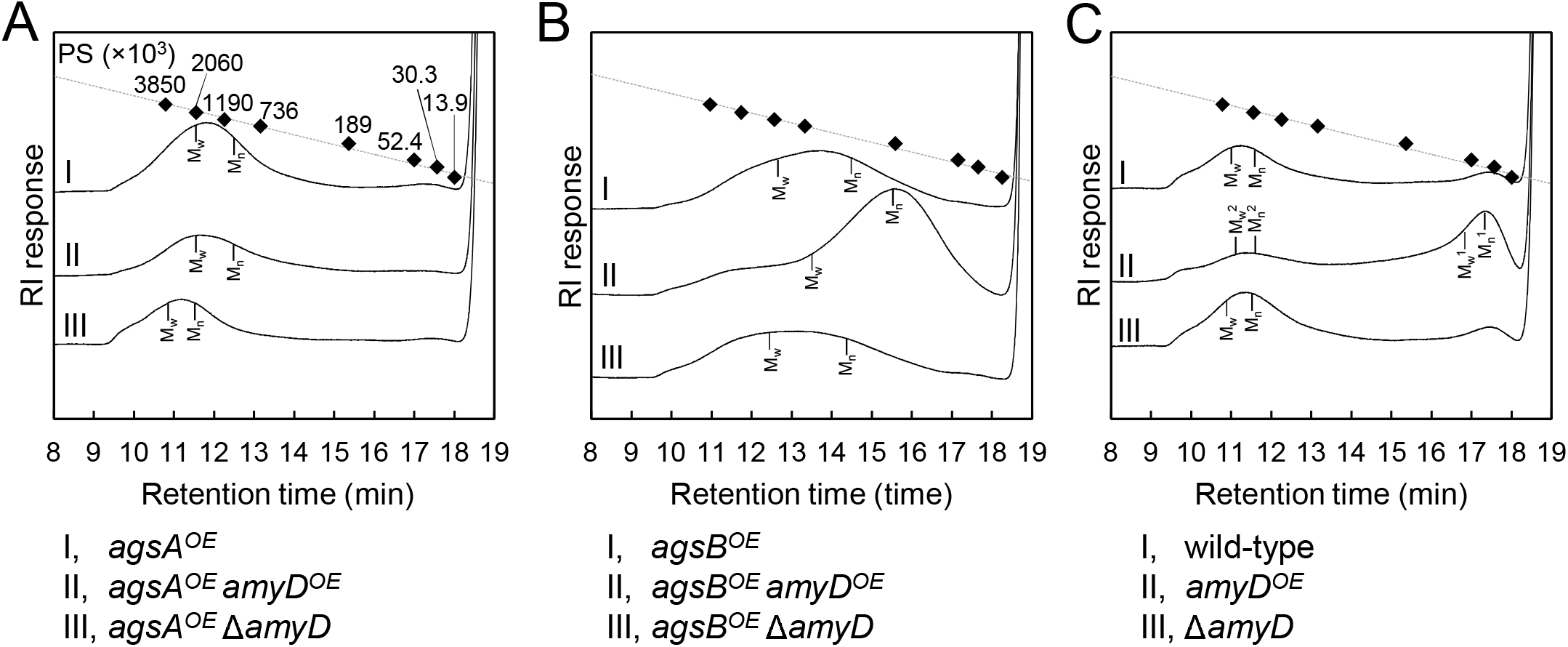
GPC elution profile of the AS2 fraction from the series of (A) *agsA^OE^* strains, (B) *agsB^OE^* strains, and (C) wild type. The AS2 fraction from 24-h-cultured mycelia of each strain was dissolved in 20 mM LiCl/DMAc. The elution profile was monitored by a refractive index detector. Molecular masses (MM) of the glucan peaks were determined from a calibration curve of polystyrene (PS) standards (♦). *M_w_*, weight-average MM; *M_n_*, number average MM.

**Table 2.**
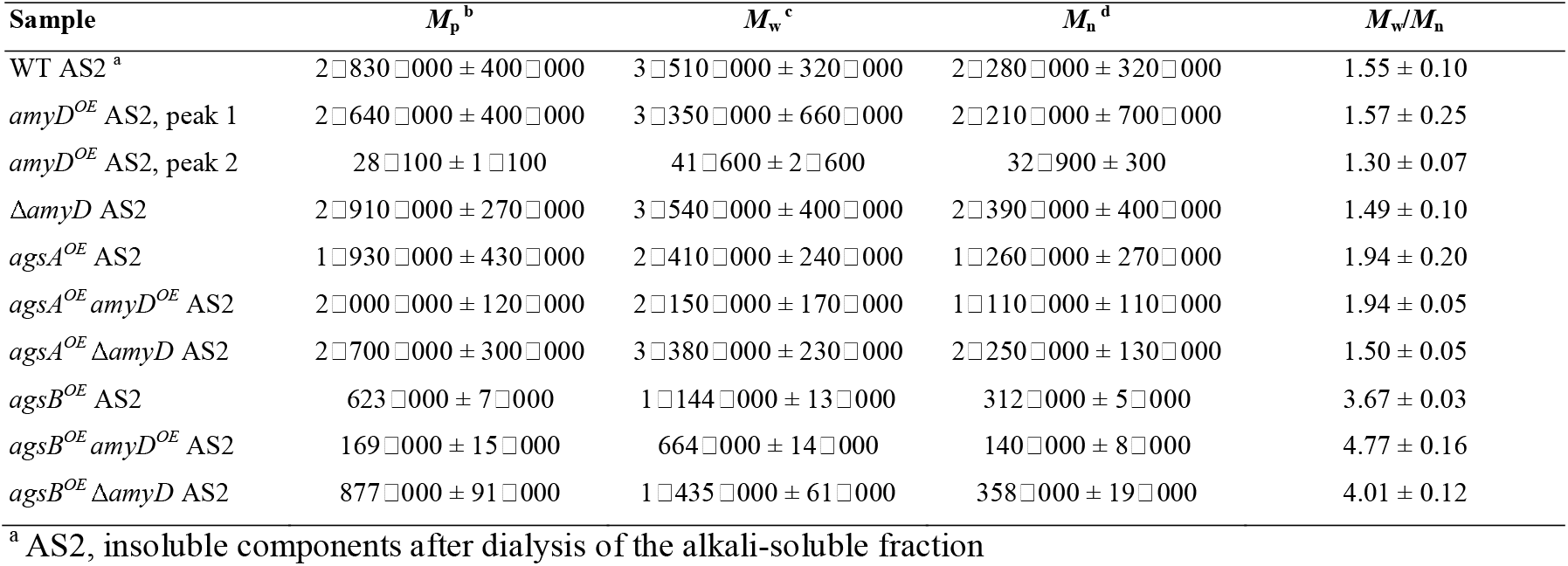
Molecular mass of alkali-soluble glucan in the cell wall.

We predicted that an unknown modification enzyme may increase the MM of alkali-soluble glucan in the endogenous *agsB*-expressing strain because the alkali-soluble glucan in these strains was synthesized mainly by AgsB. Therefore, we determined the MM of alkali-soluble glucan extracted from 16-h cultured mycelia, which should be less affected by the modification enzyme than the 24-h cultured mycelia (He et al., 2017). Unexpectedly, the *M*_n_ of the alkali-soluble glucan in the mycelia cultured for 16 h was 1 □ 980 □ 000 ± 320 000, which was similar to that in the mycelia cultured for 24 h (1 □ 930 □ 000 ± 280 □ 000; Fig. S6; Table S2). We then evaluated the MM of alkali-soluble glucan in A4, which had *M*_n_ = 2 □ 224 □ 000 ± 390□000, similar to that in the wild-type strain (Table S3).

To validate whether the degree of polymerization of α-1,3-glucan subunits in the alkali-soluble glucan was altered when the MM was changed by *amyD* disruption or overexpression, we applied Smith degradation to the alkali-soluble glucan from each strain to selectively cleave 1,4-linked glucan, and then determined the MM by GPC. One subunit of α-1,3-glucan in the alkali-soluble glucan is composed of ≈200 glucose residues (Choma et al., 2013;Miyazawa et al., 2018). The Smith-degraded alkali-soluble glucan in each strain had almost the same MM, equivalent to 300–400 glucose residues (Fig. S7; Table S4), which suggests that AmyD activity does not decrease the degree of polymerization of the glucose residues in each α-1,3-glucan subunit.

### Spatial localization of α-1,3-glucan in the cell wall is not affected by *amyD* disruption or overexpression

We previously revealed that spatial localization of α-1,3-glucan in the cell wall changes according to its MM (Miyazawa et al., 2018); α-1,3-glucans in *agsB^OE^* cells are localized in the outer layer in the cell wall, whereas most of those in the *agsA^OE^* cells are masked by a β-1,3-glucan layer. In this study, disruption or overexpression of *amyD* altered the MM of alkali-soluble glucan (Fig. 4; Table 2); therefore, we analyzed whether this alteration affected the spatial localization of α-1,3-glucan in the cell wall. In agreement with previous results (Miyazawa et al., 2018), the α-1,3-glucans with AGBD-GFP labeling showed clearly in the wild-type and *agsB^OE^* cells, but only weakly in *agsA^OE^* cells (Fig. 5). The Δ*amyD* and *amyD^OE^* cells were also labeled with AGBD-GFP (Fig. 5); fluorescent intensity in *amyD^OE^* was relatively low, which might be caused by a decrease in the amount of alkali-soluble glucan in the cell wall of *amyD^OE^* cells. The labeling with AGBD-GFP in *agsA^OE^ amyD^OE^* and *agsA^OE^* Δ*amyD* cells was weak, as was that in the cells of the parental *agsA^OE^* strain (Fig. 5). The *agsB^OE^* Δ*amyD* cells were clearly labeled with AGBD-GFP, as in the parental *agsB^OE^* (Fig. 5). The AGBD-GFP labeling was slightly weaker in *agsB^OE^ amyD^OE^* than in *agsB^OE^*, which might be attributable to a decrease in the amount of α-1,3-glucan. After treatment with β-1,3-glucanase, α-1,3-glucans of the hyphal cells in *agsA^OE^*, *agsA^OE^ amyD^OE^*, and *agsA^OE^* Δ*amyD* cells were clearly labeled with AGBD-GFP (Fig. S8), suggesting that these strains have α-1,3-glucan in the inner layer of the cell wall in their hyphal cells. Taken together, these findings indicate that disruption or overexpression of *amyD* gene scarcely affected the spatial localization of α-1,3-glucan in the cell wall.

**Figure 5.**
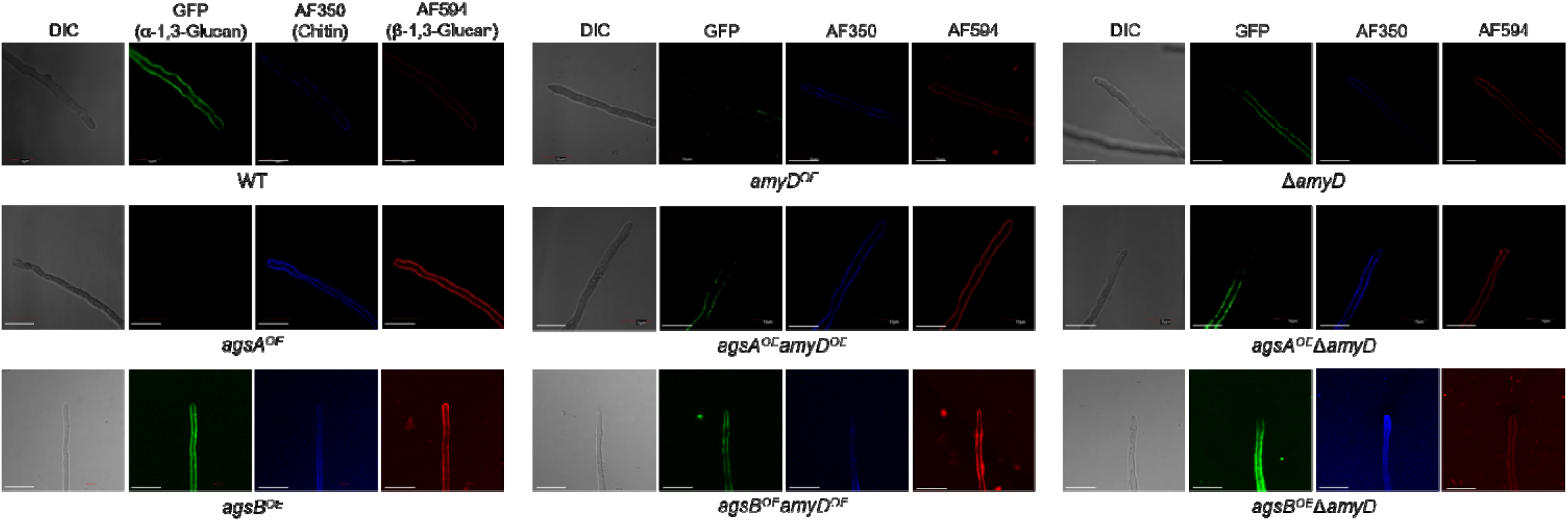
Localization of cell-wall polysaccharides of vegetative hyphae. Hyphae cultured for 16 h were fixed and stained with AGBD-GFP for α-1,3-glucan, fluorophore-labeled antibody for β-1,3-glucan, and fluorophore-labeled lectin for chitin. Scale bars are 10 μm.

### The GPI anchor is essential for the effect of AmyD on both the amount and molecular mass of alkali-soluble glucan

AmyD is thought to contain a GPI anchor at the C-terminal region. Fungal GPI anchor–type proteins are transferred from the plasma membrane to the cell wall by the activity of the GH76 family (Vogt et al., 2020). We speculated that localization in the cell wall would be essential for AmyD to reach the substrate, alkali-soluble glucan, so we constructed overexpression strains of *amyD* with and without the GPI-anchor site. Because we noticed that overexpression of *amyD* alters the phenotype or the alkali-soluble glucan, we used Δ*amyD* and *agsB^OE^* Δ*amyD* strains as hosts for the *amyD^OE^* strains. The hyphae of Δ*amyD* formed pellets in shake-flask culture (Fig. 6). Those of Δ*amyD-amyD^OE^* were dispersed, as in *amyD^OE^* (Fig. 6). Those of Δ*amyD-amyD^OE^* (ΔGPI) formed pellets, although the form was slightly different from that in the parental strain (Fig. 6). Those of *agsB^OE^* Δ*amyD*, *agsB^OE^* Δ*amyD-amyD^OE^*, and *agsB^OE^* Δ*amyD-amyD^OE^*(ΔGPI) formed similar pellets (Fig. 6). Although the Δ*amyD-amyD^OE^* hyphae had less AS2-Glc (1.13% ± 0.21%) than Δ*amyD* (5.68% ± 0.25%), the amount was restored in Δ*amyD-amyD^OE^* (ΔGPI) hyphae (5.17% ± 0.46%; Fig. 7A). These results suggest that the GPI anchor of AmyD has an important negative effect on α-1,3-glucan biosynthesis. The hyphae of *agsB^OE^* Δ*amyD-amyD^OE^* had marginally less AS2-Glc (16.2% ± 0.6%) than those of *agsB^OE^* Δ*amyD* (17.6% ± 0.3%) and *agsB^OE^* Δ*amyD-amyD^OE^*(ΔGPI) (16.7% ± 0.5%; Fig. 7B). We then evaluated the MM of alkali-soluble glucan in the cells of *agsB^OE^* Δ*amyD*, *agsB^OE^* Δ*amyD-amyD^OE^* and *agsB^OE^* Δ*amyD-amyD^OE^*(ΔGPI). The *M*_n_ of the alkali-soluble glucan in *agsB^OE^* Δ*amyD-amyD^OE^* cells (174 □ 000 ± 8 □ 000) was smaller than that in *agsB^OE^* Δ*amyD* (270 □ 000 ± 8□000; Fig. 7C; Table 3; *P* < 0.05). The *M_n_* of alkali-soluble glucan in *agsB^OE^* Δ*amyD-amy_OE_* (ΔGPI) cells (349 □ 000 ± 42 □ 000) was similar to that in *agsB^OE^* Δ*amyD* (Fig. 7C; Table 3). These results suggest that the GPI anchor of AmyD is also important for regulating the MM of alkali-soluble glucan.

**Figure 6.**
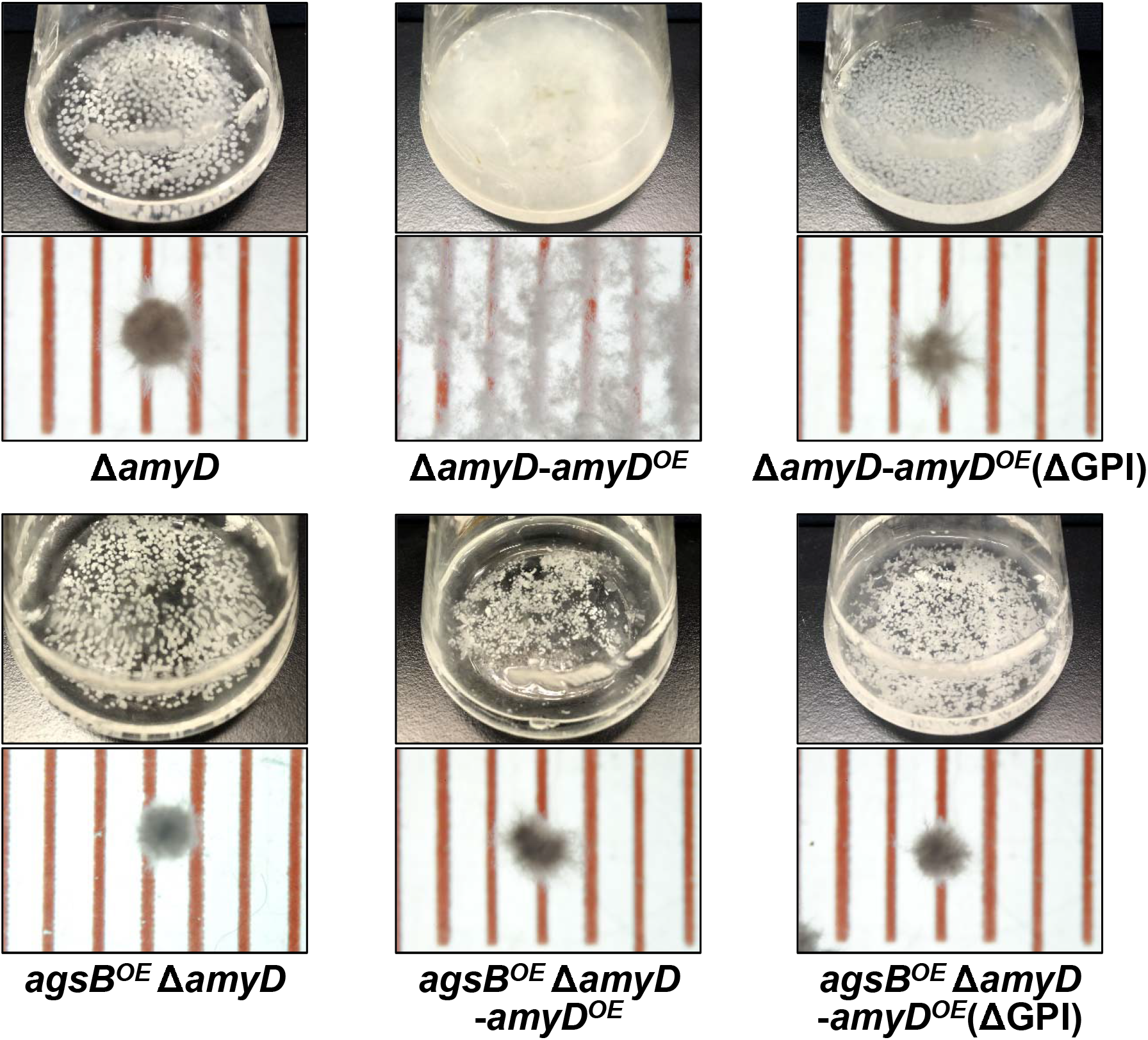
Growth characteristics of Δ*amyD-amyD^OE^* strains in liquid culture. Upper images, cultures in Erlenmeyer flasks; lower images, representative hyphal pellets of each strain under a stereomicroscope. Scale intervals are 1 mm.

**Figure 7.**
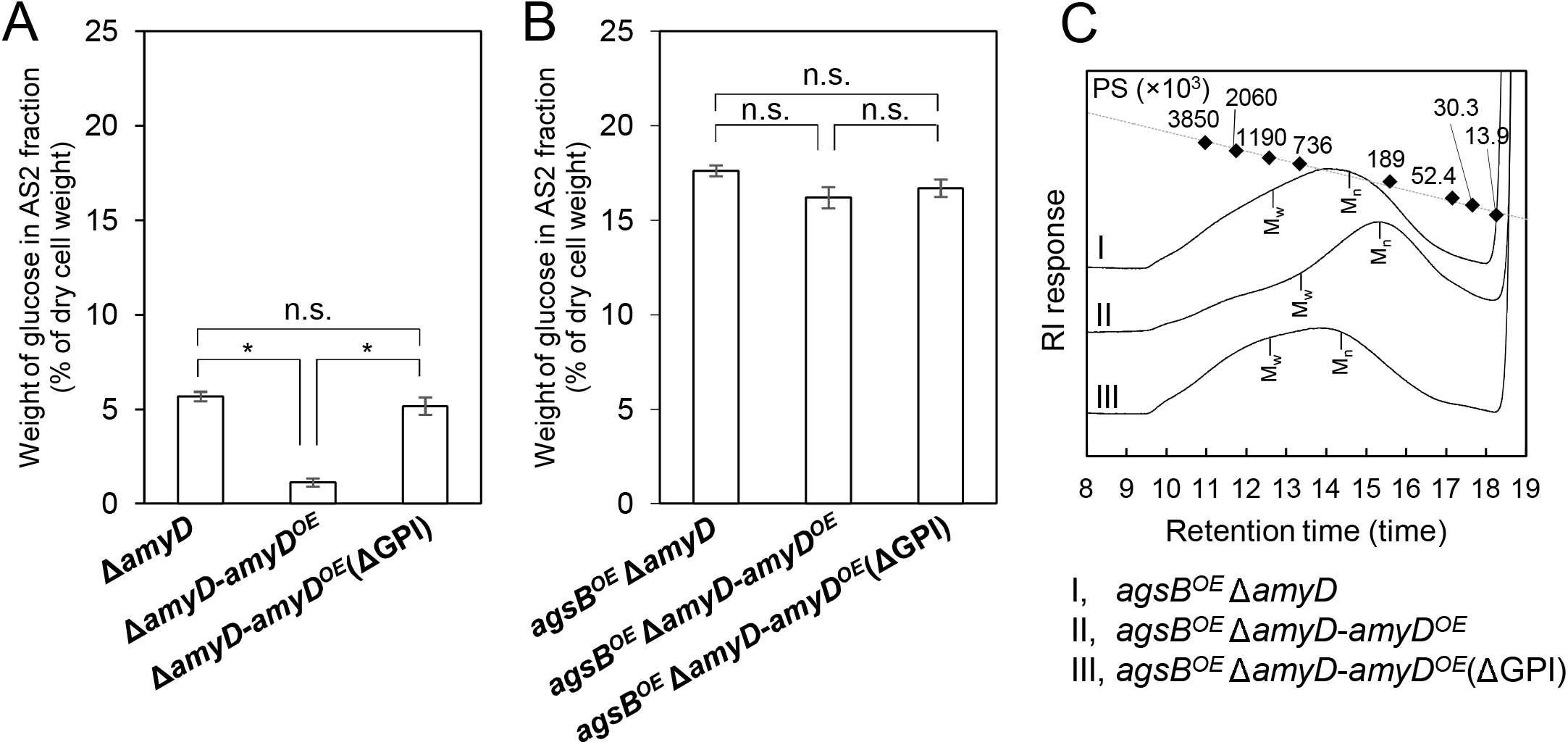
(A, B) Amounts of glucose and (C) GPC elution profiles of the AS2 fraction in Δ*amyD-amyD^OE^* strains. (**A, B**) Conidia (5.0 × 10^5^/mL) of each strain were inoculated into CD medium and rotated at 160 rpm at 37°C for 24 h. Values show glucose content of AS2 fraction as a percentage of the total cell-wall weight. Error bars represent standard error of the mean calculated from three replicates. *Significant difference by Tukey’s test (\**P* < 0.05); n.s., not significant. (**C**) The AS2 fraction from 24-h-cultured mycelia of each strain was dissolved in 20 mM LiCl/DMAc. The elution profile was monitored by a refractive index detector. Molecular masses (MM) of the glucan peaks were determined from a calibration curve of polystyrene (PS) standards (□). *M_w_*, weight-average MM; *M_n_*, number-average MM. replicates.

**Table 3.**
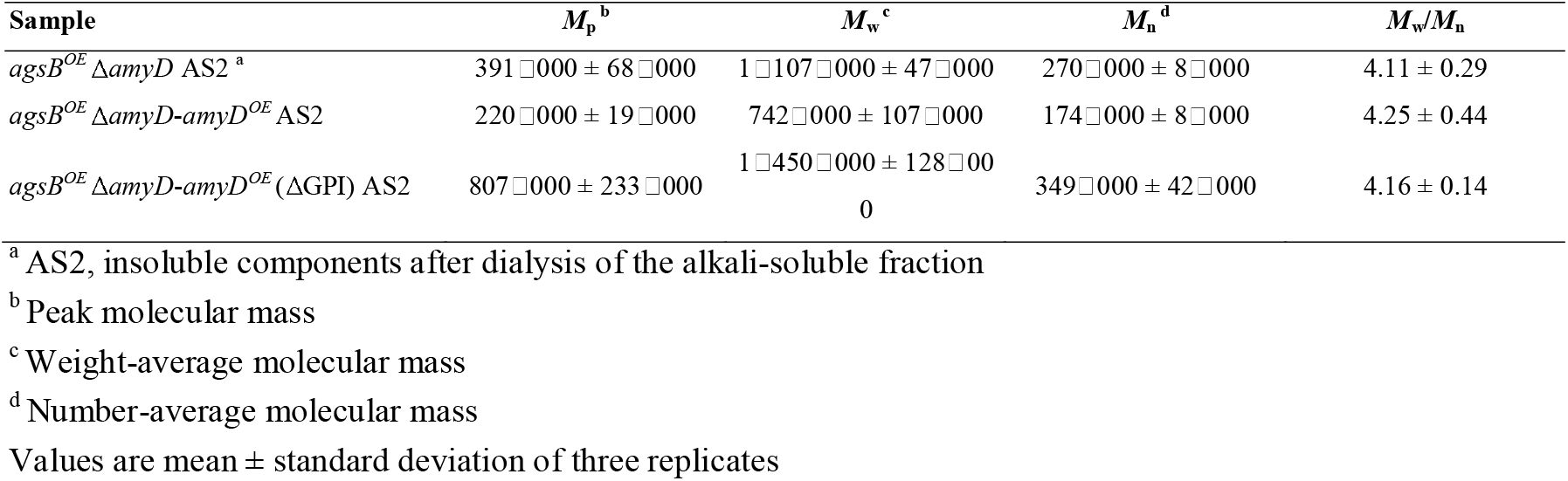
Molecular mass of alkali-soluble glucan in the cell wall of Δ*amyD-amyD* strains.

## Discussion

Although the GPI-anchored α-amylase AmyD is known to be involved in the biosynthesis of α-1,3-glucan in *A*. *nidulans* (He et al., 2014;He et al., 2017), the detailed mechanism remains unclear. Here, we looked at strains with disrupted or overexpressed *amyD* to analyze how AmyD affects the chemical properties of alkali-soluble glucan. The results reveal that overexpression of *amyD* not only decreased the MM of α-1,3-glucan, but also decreased the amount of α-1,3-glucan in the cell wall. The GPI anchor of AmyD was essential in both actions.

Overexpression of *amyD* affected the amount and MM of α-1,3-glucan in the wild-type and *agsB^OE^* strains, but not in the *agsA^OE^* strain (Fig. 3; Fig. 4; Table 2). We previously reported that the MM of α-1,3-glucan controls where α-1,3-glucan is localized in the cell wall of *A*. *nidulans;* namely, that the α-1,3-glucan with a larger MM that is synthesized by AgsA is localized in the inner layer of the cell wall, and the smaller one that is synthesized by AgsB is localized in the outer layer (Miyazawa et al., 2018). This phenomenon is explained by the fact that fungal GPI-anchored proteins are transferred from the plasma membrane to the cell wall (Orlean, 2012;Gow et al., 2017). The findings here suggest that AmyD decreased the MM of α-1,3-glucan localized at the outer layer of the cell wall. The increased MM of alkali-soluble glucan in the *agsA^OE^* Δ*amyD* strain can be explained by its GPC elution profiles which suggest that the MM of the polysaccharides was broadly distributed (Fig. 4A); in other words, *agsA^OE^* Δ*amyD* had mainly α-1,3-glucan with larger MM (>623□000 [*M_p_* of alkali-soluble glucan from *agsB^OE^*], 97.5%), but also had a small amount of α-1,3-glucan with small MM (<623□000, 2.5%). We speculate that this small amount of α-1,3-glucan with a smaller MM may be localized in the outer layer of the cell wall of *agsA^OE^*, where it is accessible to AmyD, which results in the relatively smaller MM of α-1,3-glucan. Immunoelectron microscopic analysis would be able to reveal the relationship between the spatial localization of AmyD and α-1,3-glucan in the cell wall.

AmyD of *A*. *nidulans* is considered to be a GPI-anchored protein (de Groot et al., 2009;He et al., 2014). It is well known that many fungal GPI-anchored proteins are related to remodeling of the cell wall (Samalova et al., 2020). Proteins in the “defective in filamentous growth” (DFG) family recognize the GPI core glycan and then transfer to the β-1,3- or β-1,6-glucan (Muszkieta et al., 2019;Vogt et al., 2020), which allows GPI-anchored proteins to react with their substrates in the cell wall. Although there is no direct evidence that DFG family proteins contribute to transglycosylation in *Aspergillus* species, their role in cell-wall integrity in *A*. *fumigatus* was recently reported (Li et al., 2018;Muszkieta et al., 2019), which implies that DFG family proteins are important for transferring the GPI-core glycan to β-glucan in *Aspergillus* species. To reveal the importance of the GPI anchor in the function of AmyD, we evaluated the MM and amount of α-1,3-glucan in *amyD*-overexpressing strains with or without the GPI-anchoring site. Interestingly, decreases in the MM and the amount of α-1,3-glucan were not observed when the C-terminal GPI-anchoring site was deleted (Fig. 7; Table 3); Δ*amyD-amyD^OE^* (ΔGPI) formed slightly altered pellets (Fig. 6), suggesting that AmyD expressed without its GPI anchor has only partial functions. Above all, the results show that expression of AmyD with a GPI anchor is important for reaching the substrate, α-1,3-glucan, in the space of the cell wall.

Cell-wall polysaccharides are thought to be synthesized on the plasma membrane after the secretory vesicles containing polysaccharide synthases have been exported to the hyphal tip (Riquelme, 2013). On the basis of our previous findings (Miyazawa et al., 2020), we hypothesize the process of alkali-soluble glucan biosynthesis of *A. nidulans* to be as follows: (1) the intracellular domain of α-1,3-glucan synthase polymerizes 1,3-linked α-glucan chains from UDP-glucose as a substrate from the primers, which are maltooligosaccharides produced by intracellular α-amylase AmyG; (2) the elongated glucan chain is exported to the extracellular space through the multitransmembrane domain of α-1,3-glucan synthase; (3) the extracellular domain of α-1,3-glucan connects several chains of the elongated glucan to form mature alkali-soluble glucan. The mechanism underlying the distribution of mature alkali-soluble glucan to the cell-wall network is still unknown. However, the water solubility of newly synthesized glucan might be related to the spatial distribution of α-1,3-glucan in the cell wall, because localization of α-1,3-glucan varies according to the difference in MM (Miyazawa et al., 2018). *Aspergillus niger* AgtA (encoded by an orthologue of *A. nidulans amyD*) scarcely hydrolyzes α-1,3-glucan and shows weak hydrolytic activity to starch (van der Kaaij et al., 2007). Therefore, decrease of the MM of alkali-soluble glucan in the *amyD^OE^* strain could be caused by hydrolysis of the primer/spacer residues (1,4-linked α-glucan) rather than of the 1,3-linked α-glucan region. The mechanism underlying the decrease in the amount of α-1,3-glucan by AmyD is also unknown. He et al. reported that AmyD seems to directly repress α-1,3-glucan synthesis (He et al., 2017). We suspect that AmyD with a GPI anchor on the plasma membrane binds to the spacer residues of a glucan chain that is being just synthesized by α-1,3-glucan synthase, and competitively inhibits transglycosylation by the extracellular domain of α-1,3-glucan synthase to decrease the amount of alkali-soluble glucan in the cell wall.

The *M*_n_ of the alkali-soluble glucan from the wild-type strain was larger than that from the *agsB^OE^*, although the alkali-soluble glucan from both strains seemed to be synthesized mainly by AgsB (Fig. 4; Table 2). The *M*n of the alkali-soluble glucan in the 16-h-cultured mycelia from the wild-type was similar to that from the 24-h-cultured mycelia (Fig. S6; Table S2). α-1,3-Glucan was clearly labeled with AGBD-GFP in the wild-type strain (Fig. 5). These results suggest that α-1,3-glucan was located in the outer layer of the cell wall in the wild-type strain, consistent with the localization of α-1,3-glucan synthesized by AgsB. These results imply the existence of some factor that increases the MM of α-1,3-glucan. We surmise that once a matured α-1,3-glucan molecule synthesized by AgsB is localized in the outer layer of the cell wall, macromolecules are formed by interconnecting α-1,3-glucan or connecting α-1,3-glucan to other polysaccharides, resulting in a chemically stable complex. Although the difference was not significant, the MM of Smith-degraded alkali-soluble glucan in the wildtype strain was slightly higher (Table S4) and its GPC profile had a broader distribution (Fig. S7) than those in the *agsA^OE^* and *agsB^OE^* strains, implying the existence of non-Smith-degradable glycosidic bonds (i.e. β-1,3-glycosidic bond) in the alkali-soluble fraction in the wild-type strain. It is well known that β-glucan, chitin, and galactomannan are continuously modified by hydrolase or glycosyltransferase in the cell wall (Aimanianda et al., 2017;Henry et al., 2019;Muszkieta et al., 2019). However, an enzyme that modifies α-1,3-glucan has not been reported. Recently the GPI-anchored glycosyltransferase Crh, which has a role in the crosslinking reaction for both glucan-glucan and glucan-chitin, has been reported (Fang et al., 2019). A similar enzyme that has a role in modifying α-1,3-glucan might be found soon.

Here, we revealed that AmyD in *A*. *nidulans* decreased the MM of the alkali-soluble glucan composed mainly of α-1,3-glucan in the cell wall and the amount of alkali-soluble glucan. However, a complete picture of the biosynthesis of α-1,3-glucan has yet to be described, because the substrates or proteins associated with α-1,3-glucan synthesis have not been directly demonstrated. To unveil the true nature of the biosynthesis, further biochemical analysis of the α-1,3-glucan synthase is essential.

## Author contributions

KM, AY, TN, and KA conceived and designed the experiment. KM, TY, and AT performed most experiments and analyzed the data. KM, YK, and YT performed microscopic observation. KM, AY, MS, and YY constructed fungal mutants. SK performed ^13^C NMR. AK and SY produced AGBD-GFP. KM, AY, and KA wrote the paper. KA supervised this research and acquired funding.

## Acknowledgment

We are grateful to Associate Professor Toshikazu Komoda (Miyagi University) for the NMR.

