## Supplementary material for "A glycosylphosphatidylinositol-anchored α-amylase encoded by *amyD* contributes to a decrease in the molecular mass of cell wall α-1,3-glucan in *Aspergillus nidulans*": Table S1-3, Figure S1-8

**Table S1. Primers used in this study.**

| **Purpose** | **Primer name** | **Sequence (5′ to 3′)** |
| --- | --- | --- |
| Construction of *amyD^OE^* strains | | |
|  | ANamyD-OE-LU | CCCGGGTACCGAGCTCGCGGCCAGCCAGAACGAGAC |
|  | ANamyD-OE-LL | TTCTGAGGTGCAGTTCGCGTGAGGCATGCCACATG |
|  | ANamyD-OE-RU | ATGAAAATCCTCCCATCCTTGCGCTTG |
|  | ANamyD-OE-RL | CAGTGAATTCGAGCTCTCACGCCAAAAGCAGTACG |
|  | AopyrG-PU | ACAACAGACGTACCCTGTGATGTTC |
|  | AopyrG-PL | AACTGCACCTCAGAAGAAAAGGATG |
|  | ANPtef-AopyrG-TU | GGGTACGTCTGTTGTACCAGACAGGCGCCACTC |
|  | ANPtef-ANamyD-TL | TGGGAGGATTTTCATTTTGAAGGTGGTGCGAACTTT |
| Construction of Δ*amyD* strains | | |
|  | ANamyD-D-LU | GCCATCTTTGTATTATCATTGCCACTCTC |
|  | ANamyD-D-LL | CACAGGGTACGTCTGTTGTGGCAACCTTATTGTCACCGGTTT |
|  | ANamyD-D-RU | TTCTTCTGAGGTGCAGTTACATCATCTGCAACTACAC |
|  | ANamyD-D-RL | CACCAACAATCCTTCCAC |
|  | ANamyD-D-PU | GTGACAATAAGGTTGCCACAACAGACGTACCCTGTGATGTTC |
|  | ANamyD-D-PL | GTGTAGTTGCAGATGATGTAACTGCACCTCAGAAGAAAAGGATG |
| Construction of Δ*amyD*-*amyD^OE^* strains | | |
|  | Hph-Fw | GAGGCCACTCAGGCCGATATCACC |
|  | Hph-Rv | CTGGCCTAGATGGCCGTCGACAAC |
|  | IF-Ptef1-hph-Fw | GGCCTGAGTGGCCTCACCAGACAGGCGCCACTC |
|  | IF-amyD-up-hph-Rv | GGCCATCTAGGCCAGCGCGTGAGGCATGCCACATG |
|  | ANamyD-dGPI-Fw | TGAAGCCTGAGAGCTCGAATTCACT |
|  | ANamyD-dGPI-Rv | AGCTCTCAGGCTTCACCAACAATCCT |
|  | ANamyD-IF-full-Rv | AAATTGTGCTTCTCTGCATCACGCCAAAAGCAGTACGCCCGCTAG |
|  | ANamyD-IF-DGPI-Rv | AAATTGTGCTTCTCTGCATCAGGCTTCACCAACAATCCTTCCAC |
|  | AopyrG-IF-Left-Fw | TCCCCGGGTACCGAGCTCACAACAGACGTACCCTGTGATGTTC |
|  | AopyrG-IF-Left-Rv | GACGGCCATCTAGGCCAGGGTTTCTTCGCTGAAATCGGAGAG |
|  | AopyrG-IF-Right-Fw | TGCAGAGAAGCACAATTTCCTCATC |
|  | AopyrG-IF-Right-Rv | GCCAGTGAATTCGAGCTCAACTGCACCTCAGAAGAAAAGGATG |
| PCR analysis for gene replacement | | |
|  | ANPtef-TU | AACTGCGCTTCCCAAACCAGACAGGCGCCACTC |
|  | ANagsA-tef-RL2 | CGGTACCCGGGGATCCGCGGCCGCCTGGCCAACGAAAAACATCC |
|  | ANagsA-LU1 | CGACTCTAGAGGATCCAGTGGAGGAGTTAGGGAGTGAT |
|  | ANagsA-RL1 | CGGTACCCGGGGATCCAACCGTGGTTTTGGTGGCAAAG |
|  | ANagsB-tef-RL | CGGTACCCGGGGATCCGCGGCCGCTGGCGTTAATCAGGTCGTTGG |
|  | ANamyD-up-F | AGGTTCAACGATCGAACCCAGCAAC |
|  | Hph-top-R | CCAGCTTGTGTTCCCGGTCTG |
| Quantitative PCR | | |
|  | ANHistone-RT-F | CACCCGGACACTGGTATCTC |
|  | ANHistone-RT-R | GAATACTTCGTAACGGCCTTGG |
|  | ANamyD-RT-F | GATGTCACCACCCTCGTATC |
|  | ANamyD-RT-R | GCGTAGCAATCAGCTTGTAC |

**Table S2. Molecular masses of alkali-soluble glucan in mycelia of the wild type cultured for 16 and 24 h.**

| **Sample** | ***M*_p_** **^b^** | ***M*_w_ ^c^** | ***M*_n_ ^d^** | ***M*_w_/*M*_n_** |
| --- | --- | --- | --- | --- |
| WT AS2 ^a^, 16 h | 2 270 000 ± 240 000 | 2 870 000 ± 330 000 | 1 980 000 ± 320 000 | 1.46 ± 0.08 |
| WT AS2, 24 h | 2 370 000 ± 350 000 | 2 910 000 ± 160 000 | 1 930 000 ± 280 000 | 1.53 ± 0.15 |

^a^ AS2, insoluble components after dialysis of the alkali-soluble fraction

^b^ Peak molecular mass

^c^ Weight-average molecular mass

^d^ Number-average molecular mass

Values are mean ± standard deviation of three replicates.

**Table S3. Molecular mass of alkali-soluble glucan from A4.**

| **Sample** | ***M*_p_ ^b^** | ***M*_w_ ^c^** | ***M*_n_ ^d^** | ***M*_w_/*M*_n_** |
| --- | --- | --- | --- | --- |
| A4 AS2 ^a^, 24 h | 2 240 000 ± 106 000 | 3 005 000 ± 189 000 | 2 224 000 ± 390 000 | 1.37 ± 0.16 |

^a^ AS2, insoluble components after dialysis of the alkali-soluble fraction

^b^ Peak molecular mass

^c^ Weight-average molecular mass

^d^ Number-average molecular mass

Values are mean ± standard deviation of three replicates.

**Table S4. Molecular mass of Smith-degraded alkali-soluble glucan.**

| **Sample** | ***M*_p_ ^a^** | ***M*_w_ ^b^** | ***M*_n_ ^c^** | ***M*_w_/*M*_n_** |
| --- | --- | --- | --- | --- |
| WT AS2, Smith-degraded | 72 900 ± 20 400 | 113 400 ± 29 700 | 70 100 ± 12 800 | 1.60 ± 0.14 |
| *amyD^OE^* AS2, Smith-degraded | 81 100 ± 13 300 | 138 200 ± 19 600 | 67 600 ± 2 900 | 2.04 ± 0.22 |
| Δ*amyD* AS2, Smith-degraded | 74 200 ± 24 100 | 131 700 ± 51 900 | 72 600 ± 20 100 | 1.77 ± 0.27 |
| *agsA^OE^* AS2, Smith-degraded | 57 500 ± 2 000 | 83 300 ± 7 500 | 58 100 ± 4 700 | 1.43 ± 0.02 |
| *agsA^OE^ amyD^OE^* AS2, Smith-degraded | 53 900 ± 6 300 | 70 100 ± 7 000 | 53 400 ± 5 400 | 1.31 ± 0.01 |
| *agsA^OE^* Δ*amyD* AS2, Smith-degraded | 52 300 ± 3 800 | 74 300 ± 3 300 | 54 000 ± 1,300 | 1.38 ± 0.08 |
| *agsB^OE^* AS2, Smith-degraded | 76 000 ± 5 000 | 106 000 ± 13 000 | 62 000 ± 2 000 | 1.70 ± 0.18 |
| *agsB^OE^ amyD^OE^* AS2, Smith-degraded | 68 000 ± 2 000 | 94 000 ± 3 000 | 55 000 ± 1 000 | 1.71 ± 0.06 |
| *agsB^OE^* Δ*amyD* AS2, Smith-degraded | 81 000 ± 2 000 | 130 000 ± 11 000 | 69 000 ± 1 000 | 1.88 ± 0.19 |

^a^ Peak molecular mass

^b^ Weight-average molecular mass

^c^ Number-average molecular mass

Values are mean ± standard deviation of three replicates.

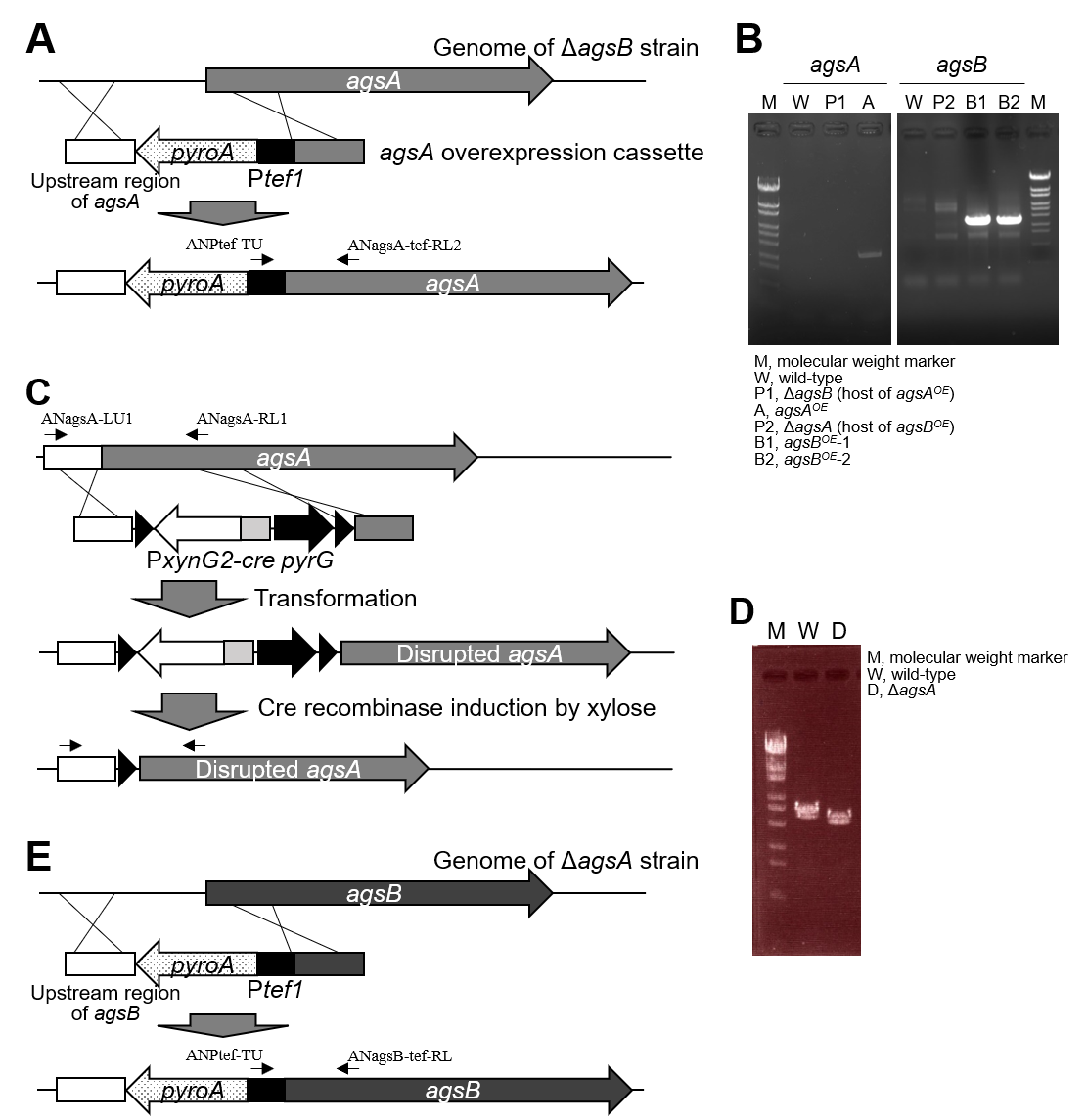

**Figure S1. Construction of *agsA*-disrupted, *agsA*-overexpressing, and *agsB*-overexpressing strains.** (**A**) Strategy for gene replacement to insert the *agsA* overexpressor cassette. (**B**) PCR analysis of *agsA* and *agsB* overexpression at the loci for *agsA* (lanes W, P1, A1, A2) and *agsB* (lanes W, P2, B1, B2) in *A*. *nidulans*. (**C**) Strategy for gene replacement to disrupt *agsA*. *Eco*RI-digested pAPG-cre/DagsA was used to transform the Δ*agsB* strain. Candidate strains were isolated by uridine and uracil requirement. Then the *Cre* gene was induced by culture on CD medium containing 1% xylose, resulting in marker cassette excision. (**D**) PCR analysis of *agsA* gene disruption at the *agsA* locus in *A*. *nidulans* using primers in **C**. (**E**) Strategy for gene replacement to insert the *agsB* overexpressor cassette.

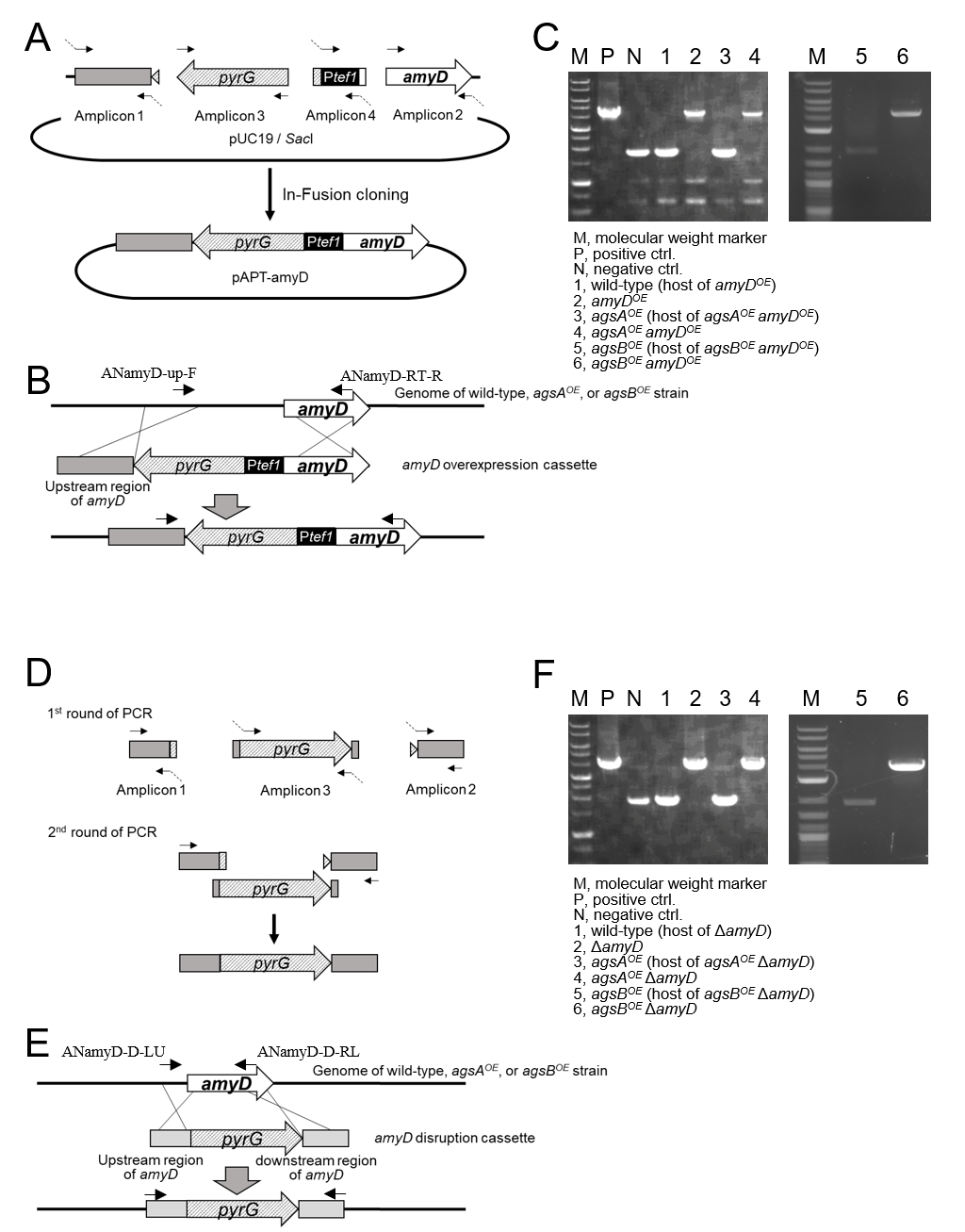

**Figure S2. Construction of *amyD^OE^* and Δ*amyD* strains.** (**A**) Construction of the pAPT-amyD plasmid for *amyD****^OE^***. Fragments containing the right and left arms of *amyD* for gene replacement, the *pyrG* marker, and the *tef1* promoter (P*tef1*) for gene overexpression were amplified and then fused to *Sac*I-digested pUC19. (**B**) Strategy for overexpression of *amyD*. *Sac*I-digested pAPT-amyD was used to transform the wild-type, *agsA^OE^*, and *agsB^OE^* strains. (**C**) PCR analysis using primers in **B** confirmed the integration of the *amyD* overexpression cassette. (**D**) Construction of the *amyD* disruption cassette. Fragments containing the right and left arms of *amyD* for gene replacement and the *pyrG* marker were amplified, and then the three fragments were fused. (**E**) Strategy for disruption of *amyD*. The fused cassette was used to transform the wild-type, *agsA^OE^*, and *agsB^OE^* strains. (**F**) PCR analysis of *amyD* gene disruption using primers in **E**.

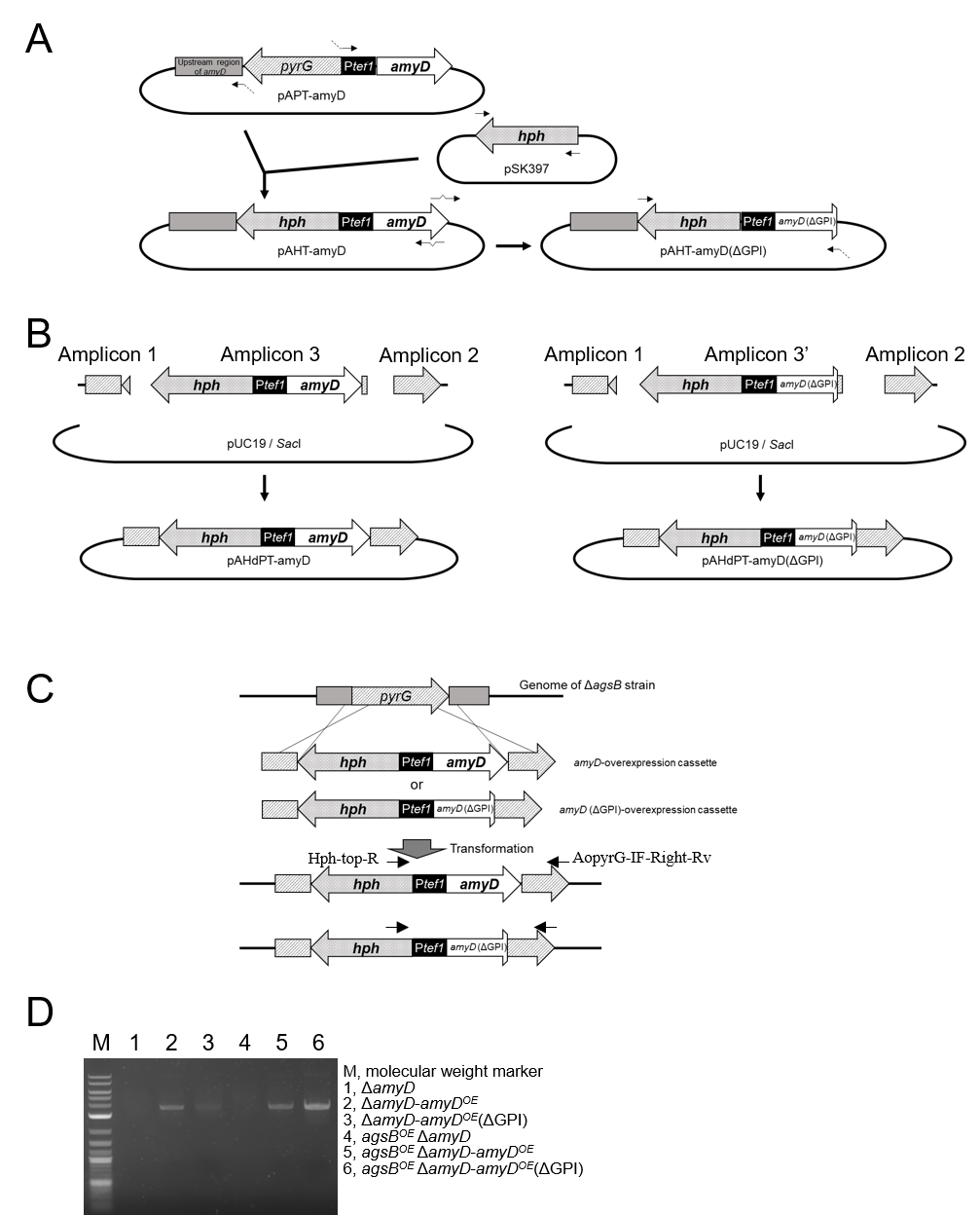

**Figure S3. Construction of complementary strains of *amyD* with and without the GPI-anchoring site.** (**A**) Construction of the plasmids pAHT-amyD and pAHT-amyD(ΔGPI). PCR amplification was performed with pAPT-amyD and pSK397 as templates, and the two fragments were fused. Then the GPI anchor–encoding region of *amyD* was deleted. (**B**) Construction of the plasmids pAHdPT-amyD and pAHdPT-amyD(ΔGPI) for overexpression of *amyD* with or without the GPI-anchor encoding region. Fragments containing the right (amplicon 1) and left (amplicon 2) arms of *pyrG* marker for gene replacement, and fragments of the *hph*, tef1 promoter (P*tef1*), and a full length (amplicon 3) or a GPI anchor region–deleted *amyD* gene (amplicon 3′) were amplified and then fused to *Sac*I-digested pUC19. (**C**) Strategy for overexpression of *amyD*. *Sac*I-digested pAHdPT-amyD and pAHdPT-amyD(ΔGPI) were used to transform the Δ*amyD* and *agsB^OE^* Δ*amyD* strains. (**D**) PCR analysis using primers in **C** confirmed the integration of the *amyD* overexpression cassette.

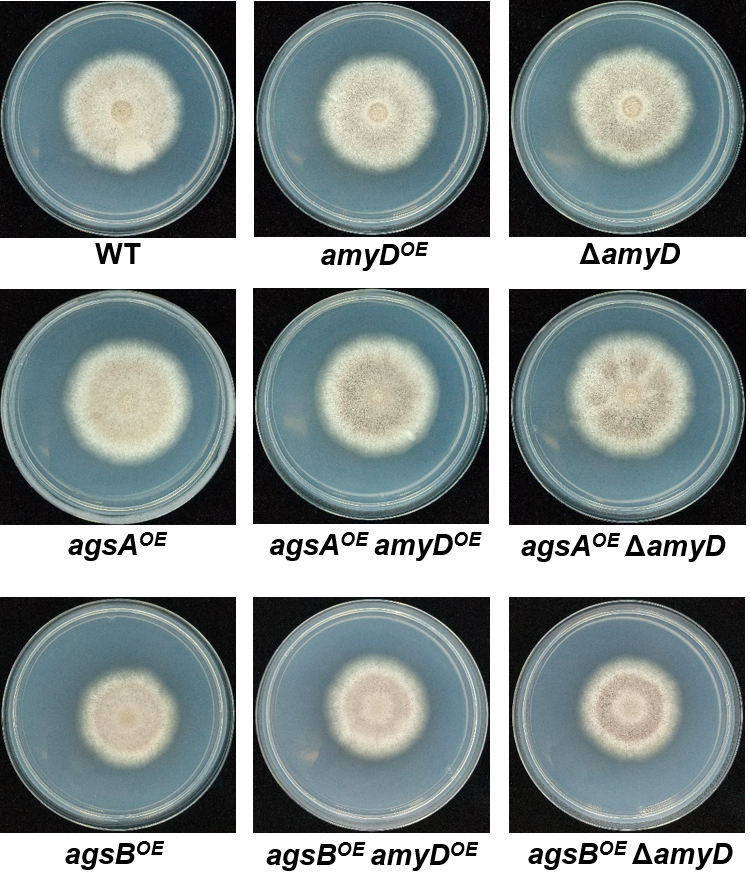

**Figure S4. Mycelial growth of the wild-type and all mutant strains on CD agar plates.** Conidia (1 × 10^4^) of each strain were inoculated at the center of a CD agar plate and incubated at 37°C for 5 days.

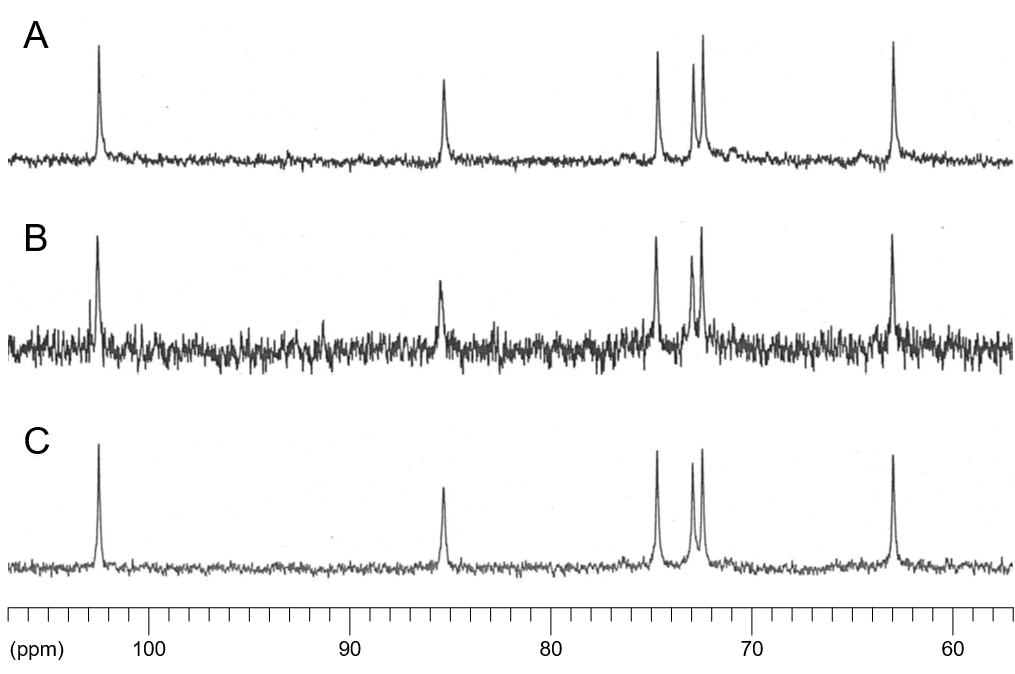

**Figure S5. ^13^C NMR spectra of the AS2 fraction from (A) wild-type, (B) *amyD^OE^*, and (C) Δ*amyD* strains.** The AS2 fraction was dissolved in 1 M NaOH/D_2_O. NMR spectra were measured at 100 MHz at 35°C. Chemical shifts were recorded relative to the resonance of DMSO-d_­6_.

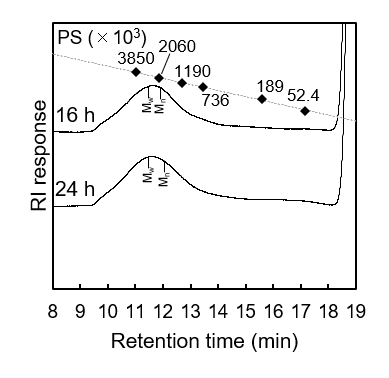

**Figure S6. GPC elution profiles of the AS2 fraction from mycelia cultured for 16 or 24 h in the series of wild-type strains.** Conidia (1.0 × 10^6^/mL) of each strain were inoculated into CD medium and rotated at 160 rpm at 37°C for 16 or 24 h. The AS2 fractions were dissolved in 20 mM LiCl/DMAc. The elution profile was monitored by a refractive index detector. Molecular masses of the glucan peaks were determined from a calibration curve of polystyrene standards.

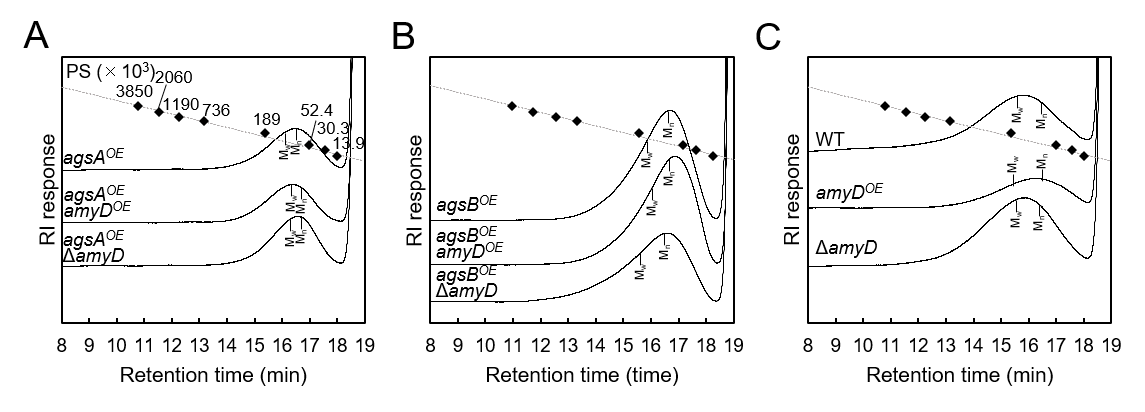

**Figure S7. GPC elution profile of the Smith-degraded AS2 fraction from the series of (A) *agsA^OE^* strains, (B) *agsB^OE^* strains, and (C) wild type.** The Smith-degraded AS2 fraction from each strain was dissolved in 20 mM LiCl/DMAc. The elution profile was monitored by a refractive index detector. Molecular masses of the glucan peaks were determined from a calibration curve of polystyrene standards.

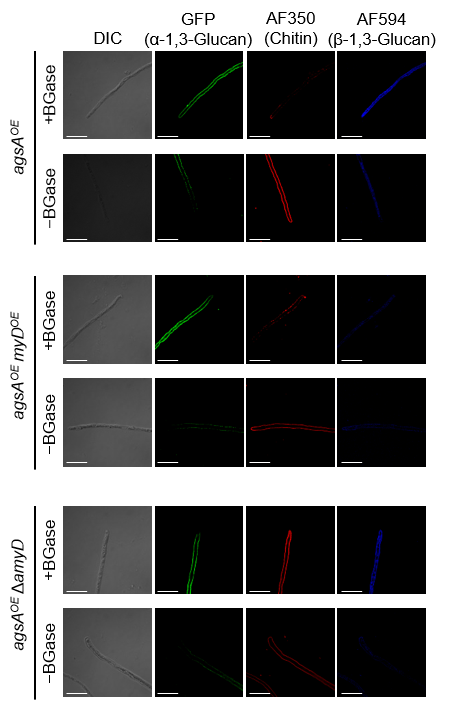

**Figure S8. Localization of cell-wall polysaccharides of the hyphal cells after treatment with β-1,3-glucanase.** Vegetative hyphae cultured for 16 h were fixed and treated with β-1,3-glucanase (BGase) for 6 h, then with AGBD-GFP for α-1,3-glucan, fluorophore-labeled antibody for β-1,3-glucan, and fluorophore-labeled lectin for chitin. Scale bars are 10 µm.
